# Collective ERK/Akt activity waves orchestrate epithelial homeostasis by driving apoptosis-induced survival

**DOI:** 10.1101/2020.06.11.145573

**Authors:** Paolo Armando Gagliardi, Maciej Dobrzyński, Marc-Antoine Jacques, Coralie Dessauges, Pascal Ender, Yannick Blum, Robert M Hughes, Andrew R. Cohen, Olivier Pertz

## Abstract

Cell death events continuously challenge epithelial barrier function, yet are crucial to eliminate old or critically damaged cells. How such apoptotic events are spatio-temporally organized to maintain epithelial homeostasis remains unclear. We observe waves of Extracellular Signal-Regulated Kinase (ERK) and AKT serine/threonine kinase (Akt) activity pulses that originate from apoptotic cells and propagate radially to healthy surrounding cells. This requires Epidermal Growth Factor Receptor (EGFR) and matrix metalloproteinase (MMP) signaling. At the single-cell level, ERK/Akt waves act as spatial survival signals that locally protect cells in the vicinity of the epithelial injury from apoptosis for a period of 3-4h. At the cell population level, ERK/Akt waves maintain epithelial homeostasis (EH) in response to mild or intense environmental insults. Disruption of this spatial signaling system results in the inability of a model epithelial tissue to ensure barrier function in response to environmental insults.

## Introduction

An epithelium is a self-organizing tissue that coordinates cell division and death to maintain its barrier function. This ability, called epithelial homeostasis (EH), is especially important due to frequent apoptosis observed in epithelia (Darwich et al., 2014). EH depends on cell density sensing, epithelial extrusion, spindle orientation, apoptotic-neighbor communication, and cell-cell adhesion dynamics to maintain epithelial integrity (Macara et al., 2014). Coordination between epithelial extrusion and cell division is crucial to maintain an adequate cell density (Eisenhoffer et al., 2012; Gudipaty et al., 2017; Marinari et al., 2012). Coordination of apoptotic cells with healthy neighboring cells is essential to close the gaps caused by extrusion of dying cells (Gagliardi et al., 2018; Gu et al., 2011).

Apoptosis can control the fate of neighboring cells. Mitogenic factors produced by apoptotic cells induce proliferation in neighboring cells, enabling wound repair (Li et al., 2010). Apoptotic cells can however also induce further apoptosis in neighboring cells during developmental processes that require coordinated cell death (Pérez-Garijo et al., 2013). Apoptosis can trigger survival fates in surrounding healthy cells in the *Drosophila* wing imaginal disk (Bilak et al., 2014). These different processes imply coordination of signaling pathways that regulate proliferation, survival or apoptosis fates. However, how these signaling pathways are regulated at the single-cell level, and spatio-temporally integrated at the population level remains poorly understood.

Mitogen-activated protein kinase (MAPK)/ERK and Phosphoinositide-3 kinase (PI3K)/Akt signaling networks are crucial for regulation of cell fate. Recent evidence has shown that ERK/Akt signaling dynamics fine-tune fate decisions at the single cell level. In epithelial cells, discrete ERK activity pulses of fixed amplitude and duration are observed at the single cell level (Albeck et al., 2013). The ERK pulse frequency depends on the epidermal growth factor (EGF) concentration, which further correlates with the efficiency of cell-cycle entry. These ERK pulses emerge from the three-tiered Raf/MEK/ERK network structure with negative feedback from ERK to Raf (Albeck et al., 2013; Ryu et al., 2015; Santos et al., 2007), that provide ultrasensitivity and adaptation properties to shape their pulsatile dynamics. In addition to ERK pulses, Akt pulses that are synchronous with the latter have been observed in epithelial cells (Sampattavanich et al., 2018). These Akt pulses are thought to maintain metabolic stability in epithelia (Hung et al., 2017). Beyond the relationship that links EGF dose and proliferation fate, little is known about how ERK/Akt regulates additional single-cell fate decisions such as survival or apoptosis, and how the latter are integrated at the cell population level to ensure EH.

Here, we show that apoptosis triggers a wave of ERK/Akt activity pulses that radially propagates for about three healthy cell layers. This signaling wave requires EGFR and MMP activity, and acts as a survival signal that locally protects cells from apoptosis for a period of 3-4 h. This single-cell behavior maintains population-level EH and tissue integrity in response to mild or intense cellular insults.

## Results

### Collective ERK/Akt activity waves propagate from apoptotic cells in quiescent unstimulated epithelium

To investigate single-cell ERK/Akt activity dynamics in epithelia, we stably transduced MCF10A cells with Histone 2B (H2B), ERK-KTR (Regot et al., 2014) and 1-396 Forkhead box O3 (FoxO3a) tagged with miRFP703 (Shcherbakova et al., 2016), mTurquoise2 (Goedhart et al., 2012) and mNeonGreen (Shaner et al., 2013) (Fig. S1A). ERK-KTR and FoxO3a biosensors respectively report single-cell ERK and Akt activity by displaying reversible nuclear-to-cytosolic translocation upon phosphorylation (Fig. S1A,B). While FoxO3a can be sensitive to ERK-dependent inputs, this has been shown to be negligible in MCF10A cells (Sampattavanich et al., 2018). We developed an automated image analysis pipeline to segment/track nuclei, and to extract cytosolic/nuclear (C/N) fluorescence intensities that quantify ERK/Akt activities (Fig. S1C). As previously shown (Albeck et al., 2013; Sampattavanich et al., 2018), starved MCF10A cells showed synchronous ERK and Akt activity pulses whose frequency increased with EGF stimulation (Fig. 1A). The amplitudes of both ERK/Akt pulses were similar with or without EGF (Fig. S1D). Visual examination (Fig. 1B), as well as spatial clustering of trajectories (Fig. 1C), revealed that starved monolayers display collective ERK/Akt activity in the form of radial waves.

**Figure 1:**
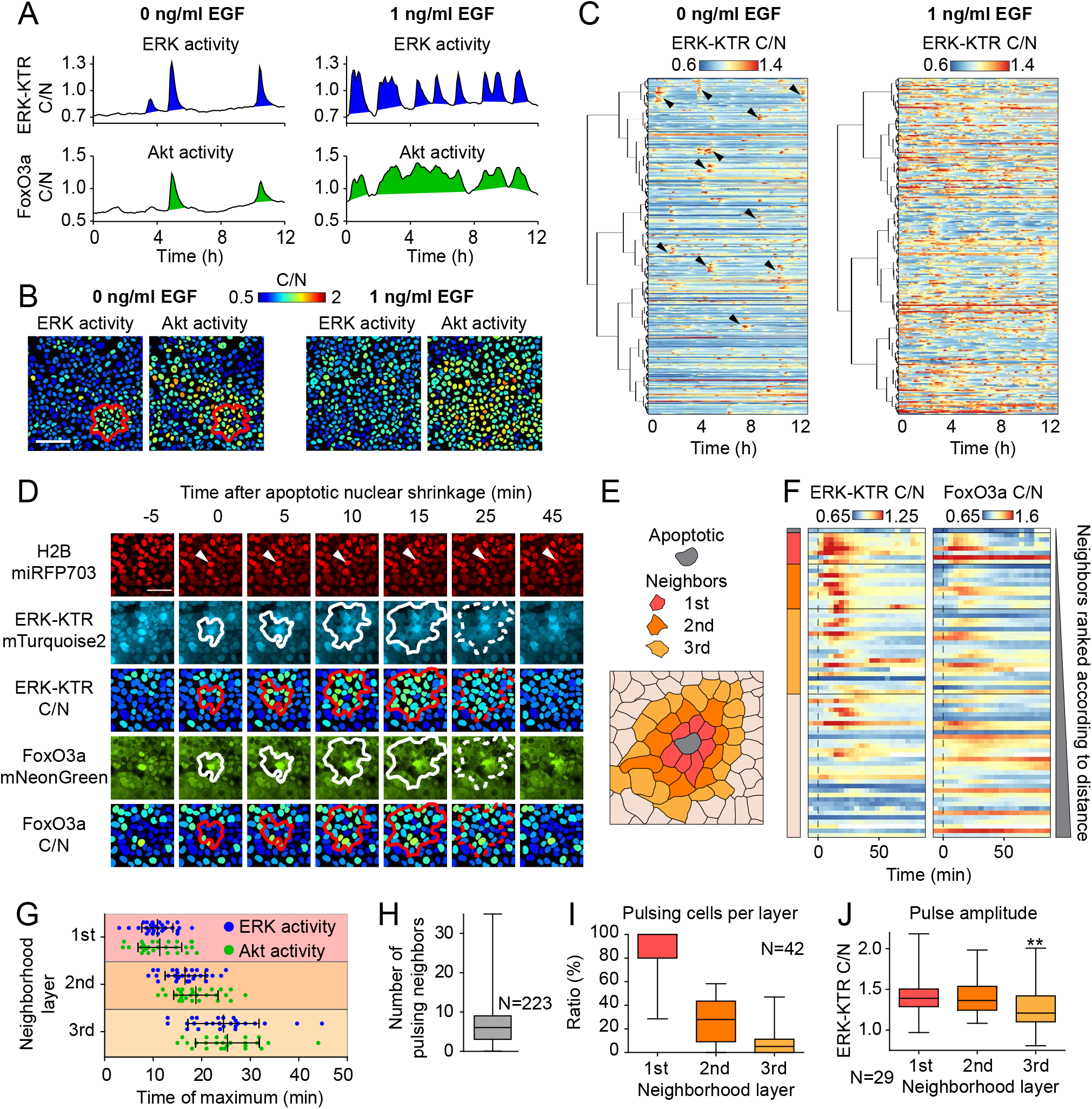
ERK/Akt waves propagate from apoptotic cells in starved MCF10A monolayers (A) ERK/Akt activity trajectories measured by cytosolic/nuclear (C/N) fluorescence ratio of ERK-KTR-mTurquoise2 and FoxO3a-mNeonGreen biosensors in starved (left) and EGF-treated (right) MCF10A monolayers. (B) Micrographs of ERK/Akt activity in MCF10A monolayers. Single nuclei are color-coded for ERK/Akt C/N ratios. The red outline delimits a collective ERK/Akt activity event. (C) Spatial clustering of ERK trajectories. Heatmaps of color-coded ERK activity trajectories clustered according to their relative time-averaged position, i.e. adjacent trajectories in the heatmap represent nearby cells. Arrowheads indicate collective ERK activity events. (D) Timelapse micrographs of one collective ERK/Akt wave originating from an apoptotic cell. H2B, raw ERK-KTR, C/N ERK-KTR, raw FoxO3a, and C/N FoxO3a images are shown. Solid outlines denote the signaling wave propagation front. Dotted outlines denote cessation of the signaling wave. Warm/cold color-coded C/N signals indicate high/low signaling activity. (E) Topology of cell outlines of the event shown in D. (F) Spatially-clustered single-cell ERK/Akt trajectories from the FOV shown in D. Trajectories are ordered according to their time-averaged distance from the apoptotic cell. Bars on the left indicate neighborhood layers from E. (G) Time of the maximum ERK/Akt activity in pulsing cells in neighbourhood layers relative to the apoptotic cell. Central vertical line is the mean, error bars are the standard deviation. (H) Distribution of the number of pulsing neighbors originating from different apoptotic events. (I) Proportion of pulsing neighbors in the neighborhood layers. (J) C/N ratio of maximal ERK-KTR amplitudes in pulsing cells in the neighborhood layers. T-test between 1st and 3rd layer (**, P < 0.01). Scale bars: (A) 100 μm, (D) 50 μm.

These ERK/Akt activity waves originated from single apoptotic cells and radially propagated in a healthy neighborhood through sequential triggering of an ERK/Akt activity pulse in each successive cell layer (Fig. 1D-F, Video S1). In the first layer, ERK/Akt activity increased synchronously with apoptotic nuclear shrinkage and peaked at 10-15 minutes after this event (Fig. 1G, S1E). In the second and third cell layers, delayed ERK/Akt activity peaked at 15-20 min and 20-30 min, respectively. An apoptotic cell could trigger ERK/Akt activity in 0 to 35 neighbors, with a median of 6 (Fig. 1H). Less than 5% of apoptotic events failed to trigger ERK/Akt (Fig. S1F). The proportion of pulsing neighbors decreased across the layers; about 90% in the 1st, 30% in the 2nd, and 10% in the 3rd (Fig. 1I). Further, ERK pulse amplitude in pulsing cells remained constant in the 1st and 2nd layer, but significantly reduced in the third layer (Fig. 1J).

These results show that apoptotic cells induce radially propagating ERK/Akt waves in neighboring healthy cells.

### ERK waves also occur in MDCK and NRK-52E monolayers, and MCF10A acini

As previously described (Aoki et al., 2013), we also observed similar ERK waves triggered by apoptosis in MDCK and NRK-52E renal epithelial cell monolayers expressing the EKAREV-NLS ERK sensor (Fig. 2A, Video S2). Analysis of signaling trajectories ranked according to the distance from the apoptotic cell showed that ERK activity propagated radially/unidirectionally in both lines (Fig. 2B). MDCK ERK waves were large, often mixed with neighboring waves, and traveled outside the field of view. NRK-52E ERK waves involved mainly 1-2 layers, leading to activation of ~20 cells (Fig. 2C,D). To exclude the possibility that signaling waves were artifacts of the monolayer culture, we evaluated

**Figure 2:**
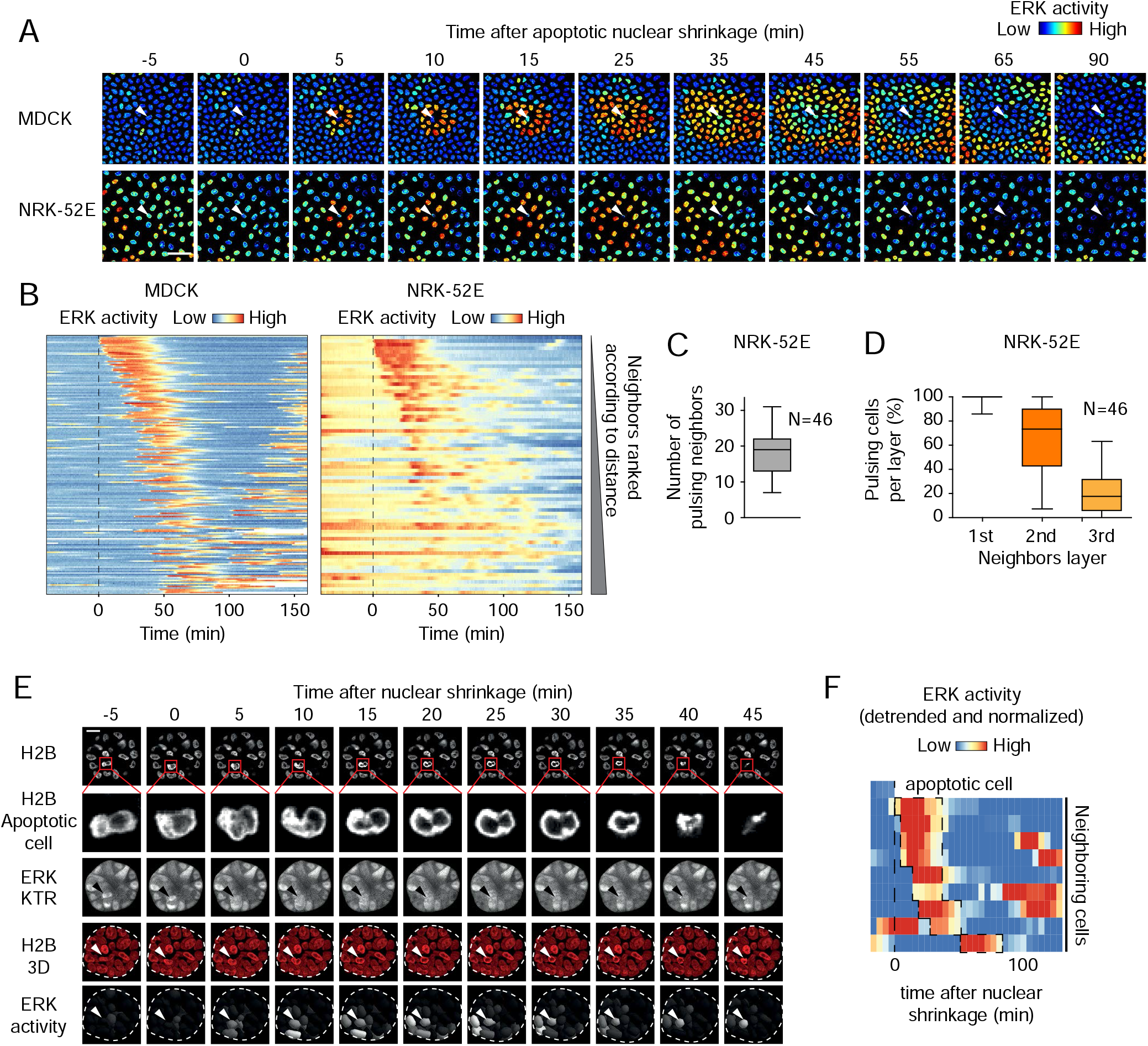
Apoptosis triggers ERK waves in MDCK and NRK-52E monolayers, and in MCF10A acini (A) Timelapse micrographs of ERK activity waves in MDCK and NRK-52E monolayers expressing the ERK activity reporter EKAREV-NLS. White arrowheads indicate the apoptotic cells. (B) Single-cell ERK/Akt trajectories ordered by the time-averaged distance from the apoptotic cell, from the events in A. Dashed lines are the time of nuclear shrinkage. (C) Distribution of the number of pulsing neighbors and (D) proportion of activated neighbors per layer in NRK-52E. (E) Timelapse micrographs and 3D rendering of an MCF10A acinus showing an apoptotic event (zoom on nucleus and arrowhead) and an ERK activity wave. The dashed line represents the spheroid contour. Denoised and normalized ERK activity is color-coded in black&white on a 3D rendering of nuclear surfaces. (F) Heatmap of denoised and normalized single-cell ERK activity trajectories of the 3D ERK wave in E (dashed line). Scale bars: (A) 50 μm, (E) 15 μm.

ERK dynamics in 3D MCF10A acini cultures (Debnath et al., 2003). We found that single apoptotic cells can trigger radially propagating ERK waves in neighboring cells (Fig. 2E,F). These waves occur alongside other, more frequent non-apoptotic ERK waves that occur spontaneously and control mammary acinar morphogenesis (Ender et al., 2020). These results show that at least ERK waves occur in different epithelial cell types, species, and culture dimensionalities.

### ERK/Akt waves are initiated during the early morphological events of apoptosis

ERK/Akt waves were triggered by apoptosis that resulted from starvation, but also when apoptosis was induced by Doxorubicin or Etoposide (Fig. S2A-C). Apoptosis is associated with a prototypical sequence of morphological events that include nuclear shrinkage, plasma membrane blebbing, chromatin condensation, cell extrusion, nuclear fragmentation and disaggregation into apoptotic bodies (Saraste and Pulkki, 2000). In epithelial cells, the two latter events are usually preceded by extrusion that removes the apoptotic body before fragmentation (Gagliardi et al., 2018; Rosenblatt et al., 2001). However, in MCF10A cells extrusion was only successful in 40% of apoptotic events, while in 60% of the events, formation of apoptotic bodies was occurring within the monolayer. The onset of the ERK/Akt waves coincided with nuclear shrinkage, the first event of the apoptosis sequence, which was then used as a reference to temporally align the collective events (Fig. 3A, G). The other morphological hallmarks of apoptosis, including epithelial extrusion, occurred after the emergence of the signaling wave, ruling them out as potential initiators of the wave (Fig. 3B-G).

**Figure 3:**
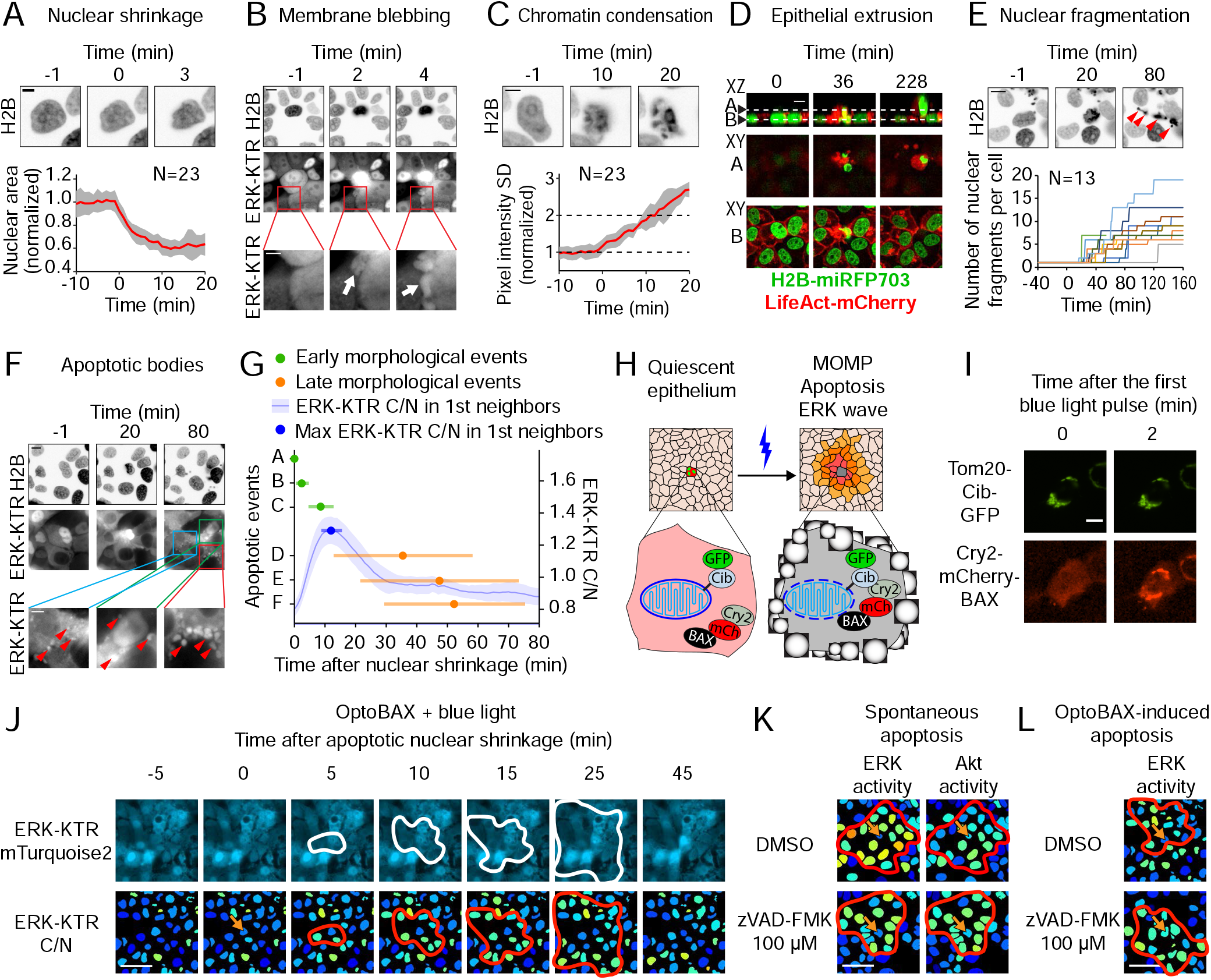
ERK/Akt waves are initiated during the early morphological events of apoptosis (A-F) Examples and quantifications of apoptotic morphological events associated with collective ERK/Akt activity waves. (A) Nuclear shrinkage. The red line and the grey area represent the mean and the standard deviation of the nuclear area, respectively. (B) Membrane blebbing. Blebs are shown in the ERK-KTR channel. (C) Chromatin condensation. Standard deviation of pixel intensity was used as a proxy of nuclear chromatin granularity. The red line and the grey area represent the mean and the standard deviation in multiple apoptotic events. (D) Epithelial extrusion. Shown are optical XZ and XY sections acquired with a spinning disk confocal microscope. (E) Nuclear fragmentation. Each line represents the number of nuclear fragments for a single apoptotic event. (F) Apoptotic bodies formation. Red arrowheads indicate individual apoptotic bodies in the ERK-KTR channel. (G) Temporal distribution of early and late morphological apoptotic events and ERK activity in the 1st layer of neighbors. Letters on the left Y-axis correspond to the morphological events shown in panels A-F. Mean and the standard deviation are shown. Blue line depicts the average ERK activity with 95% confidence interval (shade) in the first layer of neighbors. (H) Cartoon representing the OptoBAX experiment. Upon whole-field blue light illumination, OptoBAX associates to the mitochondrial membrane causing MOMP-dependent apoptosis in the central cell. (I) OptoBAX association at the outer mitochondrial membrane in a cell exposed to blue light. (J) ERK activity wave from an OptoBAX-induced apoptotic event. (K) ERK/Akt activity wave with the pan-caspase inhibitor zVAD-FMK. (L) ERK activity wave caused by Opto-BAX induced apoptosis with zVAD-FMK. Scale bars: (A, C) 5 μm, (B, F) 10 and 5 μm, (D, E, I) 10 μm, (J-L) 50 μm.

To causally link apoptosis with signaling wave initiation, we used OptoBAX, an optogenetic actuator that selectively induces apoptosis with single-cell resolution (Godwin et al., 2019). Low efficiency transfection was used to achieve stochastic expression of plasmids encoding Cry2-BAX and a mitochondrion-anchored Cib fusion in the monolayer (Fig. 3H). Upon exposure to blue light, Cry2-BAX translocates to the mitochondrion and induces mitochondrial outer membrane permeabilization (MOMP) (Fig. 3I), causing apoptosis in less than an hour. We observed that OptoBAX-induced apoptosis triggered an ERK wave identical to the one caused by spontaneous apoptosis (Fig. 3J).

To further delineate the mechanisms that trigger the signaling wave, we treated cells with zVAD-FMK, a pan-caspase inhibitor. zVAD-FMK did not prevent ERK/Akt waves in spontaneous or OptoBAX-triggered apoptosis (Fig. 3K, L, S2D-H). Further, upon zVAD-FMK treatment, apoptotic cells triggering signaling waves still displayed caspase-independent nuclear shrinkage and chromatin condensation (Fig. S2I, J) but did not exhibit caspase-dependent extrusion or apoptotic body formation (Fig. S2K). This shows that ERK/Akt waves can be triggered by MOMP and correlate with the initial morphological events of apoptosis, independently of caspase activation.

### ERK/Akt waves are triggered through EGFR and MMP signaling

To explore the signaling networks responsible for the apoptosis-triggered ERK/Akt waves, we treated starved monolayers with different inhibitors. MEK inhibition abrogated ERK waves without having any effect on Akt waves. Conversely, Akt inhibition abrogated Akt waves without any effect on ERK waves (Fig. 4A, S3A-C). This suggests a common upstream activator of ERK/Akt. Paracrine EGFR signaling initiated by MMP-mediated cleavage of pro-EGF ligands occurs in different epithelial systems (Aoki et al., 2017; Young et al., 2015). Inhibition of EGFR catalytic activity by Gefitinib, ligand binding by Cetuximab, or MMPs by Batimastat completely abrogated apoptosis-triggered ERK/Akt waves (Fig. 4A, S3D-F). OptoBax-triggered apoptosis yielded identical results (Fig. 4B, S3G). Treatment with Doramapimod and SP600125, inhibitors of p38MAPK and JNK activity respectively, showed that ERK/Akt waves are independent from these two pathways (Fig. 4A and S3I,J). Treatment of cells with Gefitinib, Batimastat or the IGF-1R inhibitor BMS754807 and simultaneous stimulation with EGF or Insulin-like GF (IGF-I), shows the specificity of EGFR inhibition on ERK/Akt activity (Fig. S4A).

**Figure 4:**
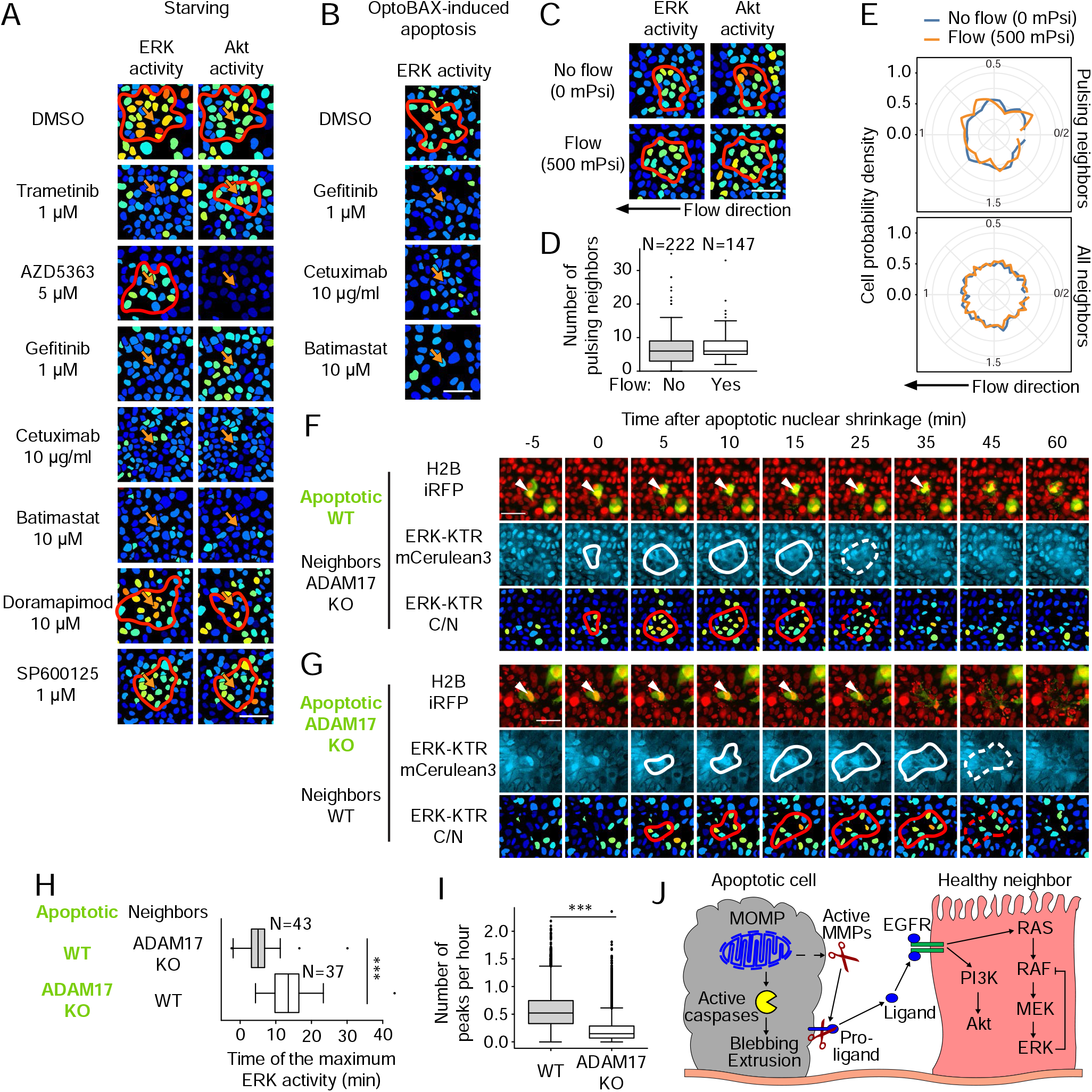
EGFR and MMP signaling mediates ERK/Akt waves (A) ERK or Akt activity waves from apoptotic cells in the presence of DMSO, MEK inhibitor Trametinib, Akt inhibitor AZD5363, EGFR inhibitor Gefitinib, anti-EGFR antibody Cetuximab, MMPs inhibitor Batimastat, p38MAPK inhibitor Doramapimod, and JNK inhibitor SP600125. Arrows indicate apoptotic cells. (B) ERK activity after OptoBAX-induced apoptosis in the presence of DMSO, Gefitinib, Cetuximab or Batimastat. (C) ERK/Akt activity waves in the absence/presence of constant flow in a microfluidic device at 500 mPsi of air pressure. (D) Distribution of the number of pulsing neighbors from different apoptotic events in the absence/presence of flow. (E) Radial distribution of neighbors that exhibit ERK pulses (top), and all neighbors (bottom) in the absence/presence of flow. (F) Timelapse micrographs of a wild-type apoptotic MCF10A cell expressing H2B-iRFP, ERK-KTR-mCerulean3 and FoxOA-mNeonGreen surrounded by ADAM17 KO cells not labeled for FoxOA-mNeonGreen. The apoptotic cell is highlighted by a white arrowhead. (G) Same as F but with a labeled ADAM17-KO apoptotic cell surrounded by unlabeled wild-type cells. (H) Distribution of the time of the maximum ERK activity of the first neighbor that shows an ERK activity pulse in several co-culture experiments like F and G. (I) ERK pulsing frequency across the entire FOV of pure wild-type vs. ADAM17 KO MCF10A. (J) Cartoon of the mechanism of the propagation of ERK/Akt activity waves. Scale bars: (A, B, C, F, G) 50 μm.

To understand whether ERK/Akt waves propagated due to free diffusion of secreted EGFR ligands, we evaluated signaling responses under flow using a microfluidic device (Ryu et al., 2015). Based on the typical diffusion coefficient of small secreted EGFR ligands (Nauman et al., 2007), we evaluated that a 25 μm/s flow was sufficient to counteract EGFR ligand’s diffusion (Fig. S4B). Using fluorescent beads, we calibrated the microfluidic device to apply the desired flow (Fig. S4C-E, Video S3). ERK/Akt waves, and their signaling trajectories did not differ in the presence or absence of flow (Fig. 4C, S4F,G). Further, the flow did not affect the size or the symmetry of waves (Fig. 4D,E), ruling out free diffusion of any EGFR ligand as a mechanism to shape waves.

Previous reports have identified ADAM metallopeptidase domain 17 (ADAM17) as the MMP involved in generation of ERK signaling waves induced by oncogenes in healthy neighbors or during wound repair in vitro (Aikin et al., 2020; Aoki et al., 2017). To explore its involvement in apoptotic ERK waves, we used ADAM17 knockout cells (ADAM17 KO) (Aikin et al., 2020). ADAM17 KO did not abrogate apoptosis-triggered ERK waves, but caused asynchronous, delayed and smaller waves (Fig. S4H-J). To understand whether apoptotic or neighboring cells require ADAM17 to transmit the signal, we performed co-culture experiments (Fig. 4F,G). Wild-type (WT) apoptotic cells within ADAM17 KO neighbors yielded normal ERK waves. In contrast, ADAM17 KO apoptotic cells within WT neighbors triggered aberrant ERK waves as described above (Fig. 4F-H), without changing the wave size (Fig. S4K). These local phenomena led to striking effects on the population-level frequency of ERK pulses. A pure ADAM17 KO cell population displayed a reduced ERK pulse frequency when compared to a WT cell population (Fig. 4I). When co-cultured, ADAM17 KO cells showed an increased pulse frequency. Conversely, WT cells showed a decreased pulse frequency (Fig. S4L,M). These results indicate MMP-mediated cleavage of pro-EGF ligands, partially through ADAM17, regulates ERK/Akt waves in an EGFR-dependent fashion (Fig. 4J).

### ERK/Akt waves are not necessary for cell extrusion

Next, we explored the function of the apoptosis-triggered ERK/Akt waves. We hypothesized that ERK/Akt waves might regulate cytoskeletal processes during epithelial extrusion (Gagliardi et al., 2018; Rosenblatt et al., 2001). This is in line with the ERK waves regulating myosin contractility during wound repair (Aoki et al., 2017). To test this hypothesis, we detected cumulated apoptotic debris extruded in the supernatant (Fig. S5A). We used starvation to trigger apoptosis and epithelial extrusion. Inhibition of extrusion caused by zVAD-FMK led to low amounts of apoptotic debris in the supernatant (Fig. S5B,C). In contrast, inhibition of ERK/Akt waves using MEK, Akt, EGFR, and MMPs inhibitors did not block this process (Fig. S5D). Direct observation of individual extrusion events in MCF10A cells (Fig. S5E) and measurements of extrusion time in MDCK cells yielded similar results (Fig. S5F). We also evaluated the assembly of the extrusion basal actomyosin ring using cells co-expressing LifeAct-mCherry and ERK-KTR-mTurquoise2 (Fig. S5G). The closure of the basal actomyosin ring occurred during the first hour after nuclear shrinkage and completely depended on caspase activity. Trametinib, Gefitinib, Cetuximab or Batimastat-mediated ERK or AZD5362-mediated Akt inhibition did not yield any noticeable effect on ring closure (Fig. S5G). Altogether, these results imply that EGFR-dependent ERK/Akt waves are not necessary for cytoskeletal processes during epithelial extrusion.

### Apoptosis-triggered ERK/Akt waves provide local survival signals

We then explored whether signaling waves regulate fate decisions. ERK pulses regulate EGF-dependent cell cycle entry (Albeck et al., 2013). However, starved MCF10A cells seldom proliferate (Ethier and Moorthy, 1991). Given that ERK/Akt signaling also regulates survival (Franke et al., 2003; Lu and Xu, 2006), we hypothesized that the signaling waves provide local survival signals. To test this, we evaluated how signaling dynamics regulates survival and apoptosis fates at the single-cell level. We restricted the analysis to ERK signaling, produced a high quality dataset of trajectories with 1’ resolution, and annotated >1000 apoptotic events. First, we compared ERK trajectories in a 6h window before apoptosis with those of non-apoptotic cells during the same window. For that purpose, we trained a convolutional neural network (CNN) to classify apoptotic and non-apoptotic cells based on ERK activity (Fig. 5A, B). A tSNE projection of the data-driven features learnt to separate the classes showed clustering of apoptotic and non-apoptotic ERK trajectories (Fig. 5A). Prototype trajectories, for which the model confidence was the highest, showed that the ERK pulse frequency is the most discriminative factor used by the CNN to separate apoptotic from non-apoptotic fates (Fig. 5B). Based on this observation, we compared ERK pulse frequency with the cell fate and with the CNN classification output. ~65% of cells that did not exhibit any ERK pulses were apoptotic. In contrast, only ~30% of cells with 2-5 pulses and 0% with >5 pulses underwent apoptosis (Fig. 5C). Evaluation of ERK trajectories in MDCK and NRK-52E cells also revealed a reduced pulse frequency in apoptotic compared to non-apoptotic cells (Fig. S6A). These results suggest that the apoptosis/survival fates in starved monolayers depend on the ERK pulse frequency.

**Figure 5:**
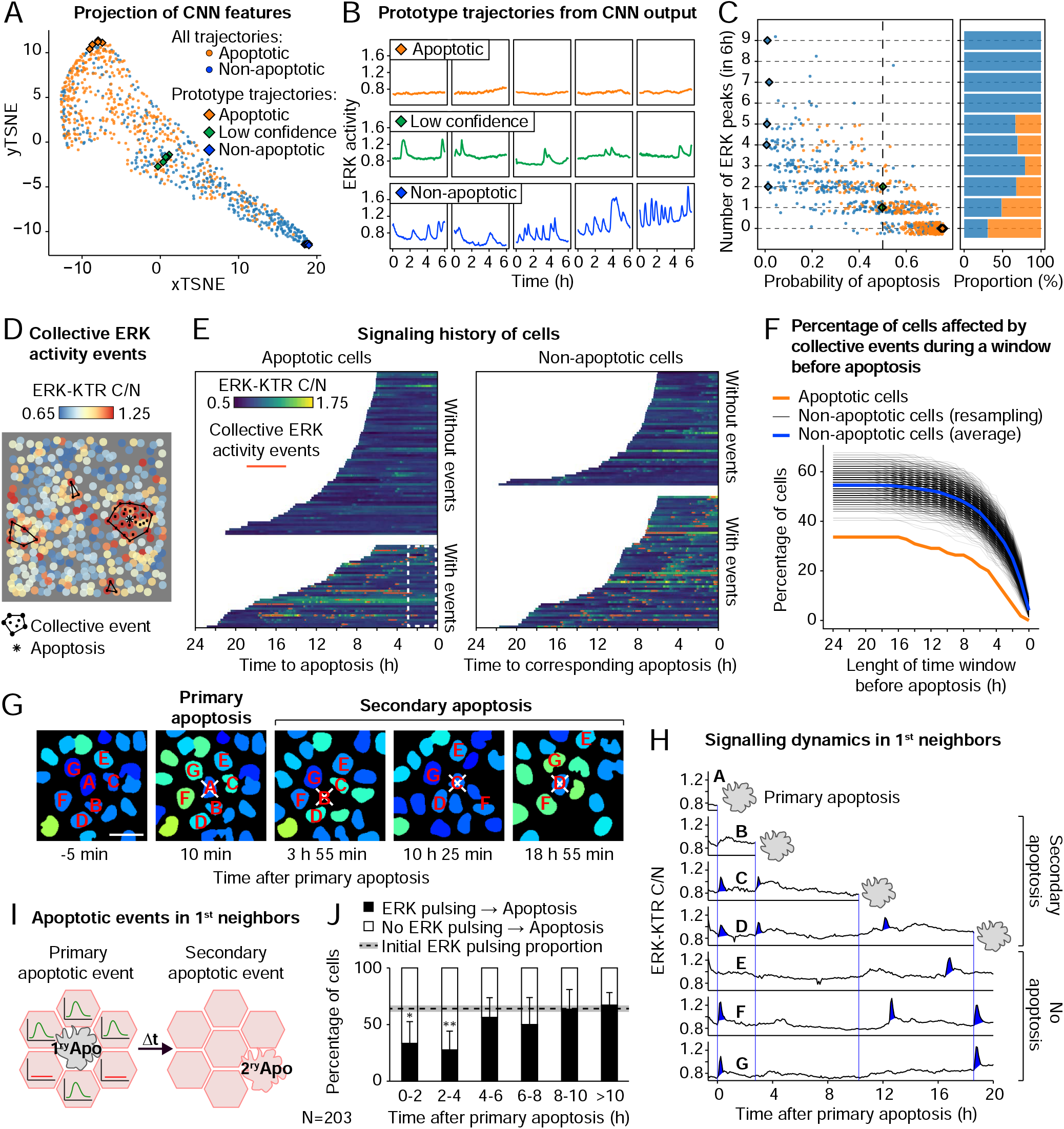
ERK waves drive AiS (A) tSNE projection of the latent features learnt by the CNN to classify ERK signaling trajectories in apoptotic vs non-apoptotic cells in the 6 h preceding apoptosis. Each dot represents a single-cell signaling trajectory. Diamonds are prototype ERK trajectories of apoptosis (orange), non-apoptosis (blue) or low-confidence prediction (green). (B) Prototype ERK trajectories from A. (C) Distribution of ERK trajectories according to the probability of apoptosis predicted by CNN against the number of ERK activity pulses in the trajectories. The right plot shows the proportion of apoptotic vs non-apoptotic cells. (D) Automatic identification of collective events based on ERK activity. Black polygons indicate collective events; asterisks, apoptosis. (E) Heatmaps of ERK activity in apoptotic vs. non-apoptotic cells. The orange overlay indicates the times when cells are involved in collective events. The white dashed box indicates a 3h window before apoptosis. (F) Percentage of cells affected by collective events during a window before apoptosis. The orange line corresponds to apoptotic cells. Black lines represent the result from paired non-apoptotic cells, resampled 1000 times in the same field of view as the corresponding apoptotic cells. Blue line is the average of all resampled curves. (G) Example of ERK activity dynamics in consecutive apoptotic events (white crosses) in the first layer of neighbors of a primary apoptotic event indicated “A”. Scale bar, 20 μm. (H) ERK signaling trajectories from G. (I) Classification of 1st neighbors of apoptotic cells according to ERK pulsing. (J) Percentage of secondary apoptotic events in the 1st neighbors that received an ERK activity pulse during 2h intervals after the primary apoptotic event. Error bars represent 95% confidence interval. Dashed line and shaded grey area represent the percentage of pulsing cells in all 1st neighbors and 95% confidence interval. Significance with respect to “>10h” calculated with a Chi-square test and corrected with the Holm-Bonferroni method (*, P < 0.05; **, P < 0.01).

Second, we quantified differences in participation of apoptotic and non-apoptotic cells in collective signaling events. We developed a computational approach that automatically identifies collective ERK waves (Fig. 5D, S6B). We found that 24% (SD=6) of ERK pulses were part of such events and that 67% (SD=8) of collective waves were triggered by apoptosis within a 20 min time window. Then, we temporally aligned ERK trajectories of apoptotic cells with respect to apoptotic events, overlaid occurences of collective events, and compared the incidence of the latter with those of non-apoptotic cells (Fig. 5E). Collective ERK signaling events occurred in only ~32% of apoptotic trajectories, compared to ~53% non-apoptotic cells (Fig. 5E, F). Moreover, ERK pulsing during a 3h window before apoptosis was less frequent than in the time before (Fig. 5E and S6C). These results indicate that cells that experience collective ERK events are less likely to die than cells that do not.

Third, we explored whether an ERK pulse within an apoptosis-triggered signaling wave locally promotes survival. Thus, we evaluated the signaling history of “secondary” apoptotic cells located within the 1st layer of neighbors of a primary apoptotic event (Fig. 5G-I, Video S4). We found that up to 4h after the primary apoptosis, secondary apoptosis is significantly less likely to occur in cells that experienced an ERK pulse induced by the primary event (Fig. 5J). We obtained the same result with the 2nd layer of neighbors (Fig. S6D). Evaluating the cumulative distribution of the probability of a secondary apoptotic event revealed a lower apoptosis probability for neighbors that received a pulse from the primary apoptotic event compared to those that did not (Fig. S6E). This strongly suggests that apoptosis induces waves of ERK and Akt activity that locally induce survival for approximately 4h. We term this process apoptosis-induced survival (AiS).

### ERK pulse frequency determines survival fate

To establish whether specific signaling dynamics regulates the survival fate, we used 2 optogenetic systems to evoke different signaling pulse frequencies. We simultaneously measured ERK activity using spectrally-compatible ERK-KTR-mRuby2 (Fig. 6A). The first system is a photo-excitable fibroblast growth factor 1-based receptor (OptoFGFR) that activates both ERK and Akt signaling (Kim et al., 2014), and thus mimics the EGFR-dependent signaling observed in our cell system. The second is a photo-excitable RAF construct (OptoRAF) that selectively controls ERK activity (Aoki et al., 2017). For both systems, a 100 ms of 3 W/cm^2^ pulse of blue light could induce a robust ERK pulse of similar amplitude (Fig. 6B, C, Video S5). We induced ERK pulses with light stimulation applied every 1, 2, 3, 4, 6, 12 h with both optogenetic systems during a 24h period after starvation (Fig. 6D, E and G). We measured a robust reduction of the apoptotic rate when ERK pulses were triggered at least every 3h with OptoFGFR (Fig. 6F, Video S6). At the cell population level, this almost completely suppressed the peak of apoptosis triggered by starvation (Fig. S6F). ERK or Akt inhibition abrogated the protection granted by the stimulation of OptoFGFR at high frequency (Fig. S6G). We observed similar effects with OptoRAF in which high stimulation frequencies (1, 2, 3, 4h pulse periodicity) provided robust protection, which then gradually diminished at lower frequencies (Fig. 6H). Contrary to OptoFGFR, the effect of OptoRAF pulses on survival was independent from Akt activity (Fig. S6H) and completely dependent on the MAPK pathway (Fig. S6I-J). Using Batimastat, that unspecifically increases the apoptosis rate regardless of light pulses, we also excluded a possible ERK-ADAM17-EGFR axis through which OptoRAF could exert its pro-survival effect (Fig. S6K). Using both optogenetic systems, ERK pulses induced every 3-4h were sufficient for survival. OptoFGFR mediated its effect through both ERK and Akt, while OptoRAF only through ERK. However, the expression of the OptoRAF system increases basal survival (compare Fig. 5F and 5H), introducing a small bias in our system.

**Figure 6:**
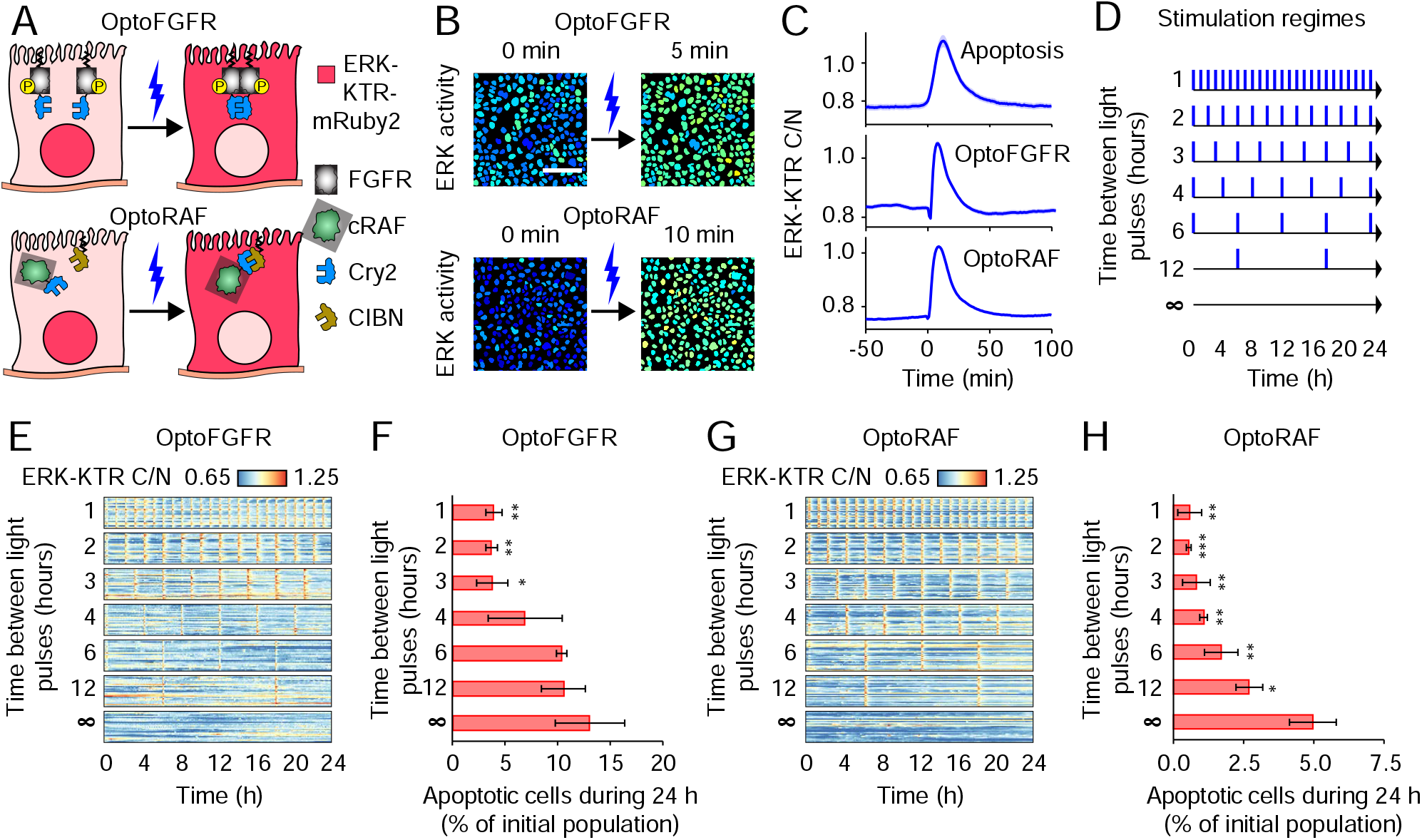
Frequency of ERK pulses determines the survival fate (A) Cartoon of the two optogenetic tools used to induce synthetic pulsed ERK signaling regimes. OptoFGFR is a CRY2-based, blue-light-dimerizable, plasma-membrane linked FGFR1 intracellular domain. OptoRAF consists of cRAF fused to CRY2 and a plasma membrane-targeted CIBN domain. (B) Representative micrographs of ERK-KTR C/N ratio in OptoFGFR and optoRAF expressing cells before and after a blue light pulse. Scale bar 100 μm. (C) Average ERK activity trajectories from apoptotic neighbors, or from cells expressing OptoFGFR or OptoRAF in response to blue light. Mean and 95% confidence interval are shown. (D) Different blue light stimulation regimes ranging from 1 pulse/hour to no pulsing. (E) ERK signaling trajectories in OptoFGFR-expressing cells responding to the stimulation regimes in D. (F) Apoptotic rate of OptoFGFR-expressing cells during 24h after starvation in response to the stimulation regimes in D. (G,H) Same as panels E and F but using OptoRAF-expressing cells. Error bars represent the standard deviation of 3 replicates. T-test with respect to unstimulated cells (*, P < 0.05; **, P < 0.01; ***, P < 0.001).

### AiS maintains epithelial integrity in response to environmental insults of different intensities

AiS might contribute to population level EH and epithelial integrity in response to environmental insults. To explore that, we evaluated the rate of apoptosis in response to starvation and compared it with population-averaged ERK/Akt activity (Fig. 7A). Monolayer starvation resulted in a transient peak of apoptosis that started 2-3h after starvation, and lasted for another 4-5h until a steady-state, low apoptosis rate ensued. Strikingly, population-averaged ERK/Akt activities immediately decreased with starvation and transiently reactivated with kinetics that were slightly delayed with respect to the apoptotic rate. This suggests that the whole monolayer can adapt to starvation-induced stress by dynamically regulating survival to maintain EH and epithelial integrity.

**Figure 7:**
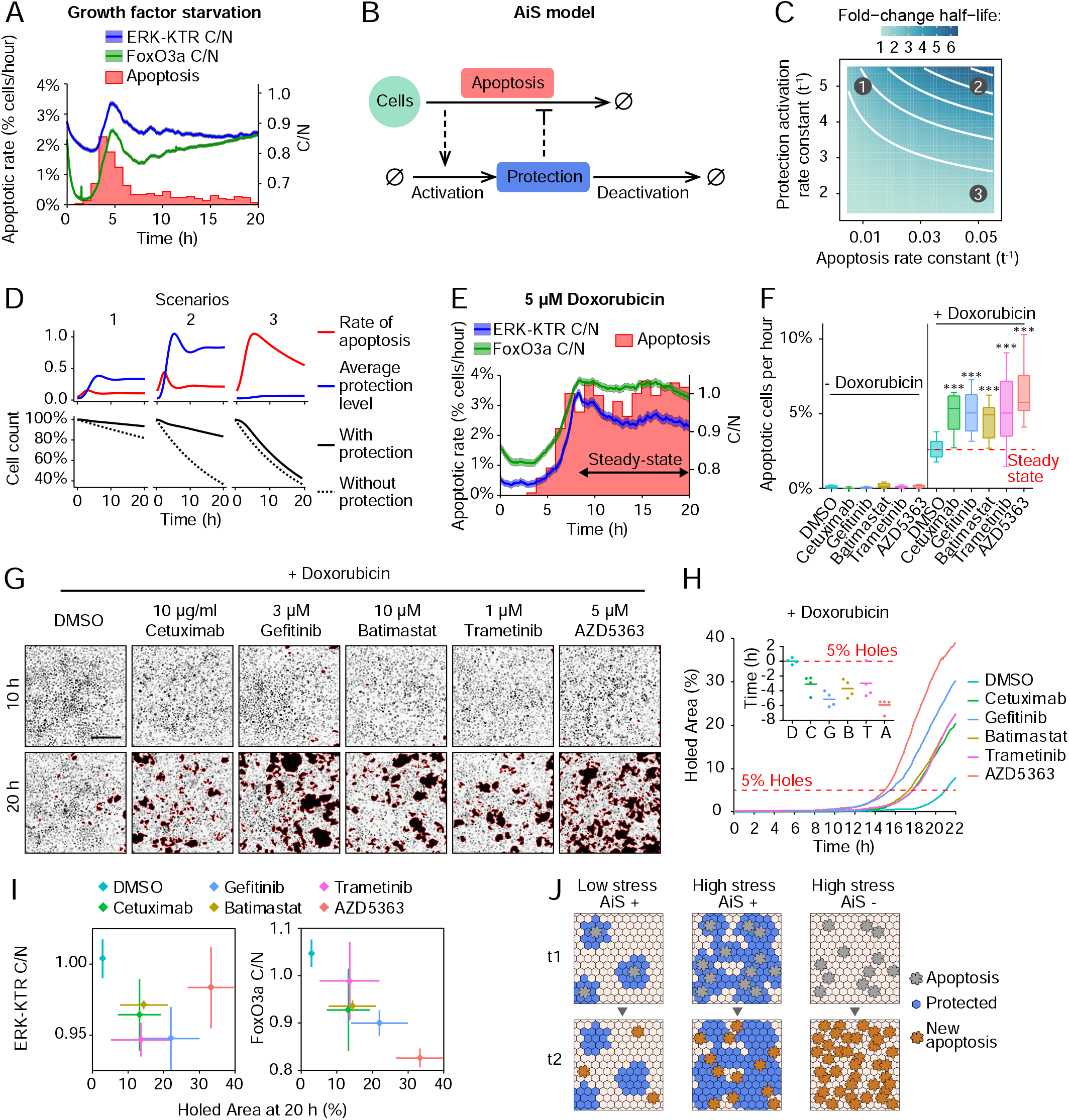
AiS maintains epithelial tissue integrity (A) Rate of apoptotic cells per hour after starvation, superimposed with average ERK/Akt activity trajectories of all cells in the analyzed FOVs. Shaded areas represent 95% CI. (B) Two-component model of AiS. The rate of apoptosis induces the protection, which then inhibits apoptosis in the neighbors. The onset of protection is multi-stage, which introduces a delay in the apoptosis inhibition. (C) Exploration of the model parameter space. Half-decay time of the initial pool of cells calculated as a fold-change with respect to the half-decay time of an exponential decay without induction of protection. Isolines represent 2-6 fold-change half-life. Points indicate 3 scenarios explored in the next panel. (D) Predicted rate of apoptosis, protection level and the cell count for 3 scenarios corresponding to: 1 - starved untreated epithelium, 2 - acute pro-apoptotic treatment, 3 - acute pro-apoptotic treatment and AiS inhibition. The cell count is compared to an exponential decay without protection. (E) Rate of apoptotic cells per hour after treatment with 5 μM Doxorubicin, superimposed with average ERK and Akt activity trajectories. Shaded areas represent 95% CI. (F) Distribution of the rate of apoptotic cells per hour 8-20h after Doxorubicin addition and in presence of different treatments. (G) Micrographs of the nuclear channel superimposed with holes detection. Scale bar 200 μm. (H) Percentage of holed area from G. Each line is the average of 4 different fields of view. 5% of area occupied by holes has been used as a reference to generate the inset chart. Each dot represents a field of view and the horizontal lines the average value. D, DMSO; C, Cetuximab; G, Gefitinib; B, Batimastat; T, Trametinib; A, AZD5363. (I) Correlation between ERK or Akt activity and the percentage of the holed area 20h after Doxorubicin addition. Diamonds, average; error bars, standard deviation. (J) Cartoon of the population-scale effect of AiS on the epithelium exposed to low or high apoptotic stress in comparison with the same epithelium that lacks AiS.

We built a mathematical model to capture the dynamic relationship between apoptosis and survival. The model consists of 2 interacting components: apoptotic cells, and the protection from cell death mediated by ERK/Akt activity (Fig. 7B). The increase in protection level depends on the rate of apoptosis, which in turn is negatively regulated by the amount of protection in the system. We varied the rate constants of apoptosis and protection activation, and calculated the fold change in the half-life of the initial cell pool with respect to the half-life of a pure exponential decay, i.e. a model without the protection component (Fig. 7C). We investigated 3 scenarios (Fig. 7C, D). In scenario 1, we set a low rate of apoptosis and a high rate of protection activation to mimic our starvation experiment (Fig. 7D). The model agreed with the starvation experiment (Fig. 7A) in that an episode of increased apoptosis is followed by a delayed induction of protection. In scenario 2, we increased the rate of apoptosis (e.g. as possibly induced by a cytotoxic drug), which yielded higher apoptosis and protection levels after the initial transient (Fig. 7D upper panel). Although the cell number decreases faster than in scenario 1, the high protection level prevents cells from dying with the rate of a pure exponential decay (Fig. 7D, lower panel). In scenario 3, we maintained the high apoptosis rate but we lowered the protection rate (e.g. inhibition of ERK/Akt activity). Here, the apoptosis rate exhibits a much slower transient due to a weaker protection (Fig. 7D upper panel), leading the cell count to closely follow a pure exponential decay (Fig. 7D, lower panel).

To experimentally validate the results obtained from the model (scenarios 2/3), we treated our cells with doxorubicin, which induces apoptosis by oxidative stress in MCF10A cells (Gajewski et al., 2007). A dose response screen identified 2.5 and 5 μM doxorubicin as concentrations that induce potent cell death (Fig. S7A). This induced a steep apoptosis increase from 5 to 7 h after incubation, followed by steady apoptosis levels that corresponded to approximately 3-4% of the cell population per hour for more than 10 h (Fig. S7B). The steep apoptosis increase was followed by a slightly delayed increase of ERK/Akt population average activity that then remained high (Fig. 7E, S7C and D). Lower doxorubicin doses that promoted low apoptosis only led to low ERK/Akt activity (Fig. S7C and D). These results are in line with the predictions of scenario 2 of our model (Fig. 7D). To exclude that ERK/Akt activation was due to accumulation of EGFR ligands in the supernatant, we kept the cells under constant perfusion (Fig. S4C). With doxorubicin, we observed an increase in the apoptotic rate reaching about 4% at 13h post addition (Fig. S7E). This also resulted in higher ERK/Akt activities compared to the untreated control (Fig. S7F-H). This excludes accumulation of ligands as the cause of the global increase in ERK/Akt activity.

To explore if this sustained population-level ERK/Akt activity was due to AiS, we inhibited EGFR, MMPs, ERK or Akt and evaluated apoptosis rates. While individual inhibitors had only modest effect on apoptosis, their combination with doxorubicin caused a dramatic increase in the apoptotic rate (Fig. 7F, S7I and J), as predicted in scenario 3 of our model (Fig. 6D). To characterize how changes in apoptosis rate impact epithelial integrity, we measured hole formation in the monolayer. We found that doxorubicin-drug combinations quickly led to loss of epithelial integrity ~15h after drug treatment, while doxorubicin alone only led to few holes (Fig. 7G and H, Video S7). Further, we observed an inverse correlation between population-averaged ERK/Akt activity levels and the holed area 20h after doxorubicin treatment (Fig. 6I). These results strongly suggest that AiS contributes to epithelial integrity by reacting to environmental insults of different intensities (Fig.6J).

## Discussion

Epithelial tissues require dynamic coordination of apoptosis, survival and proliferation to maintain a critical cell density and barrier function (Macara et al., 2014). We report on AiS, a signaling mechanism that preserves EH in response to environmental stresses (Fig. 6J). At the single-cell scale, AiS involves apoptosis-triggered ERK/Akt waves that protect nearby cells from apoptosis for 3-4h. At the population scale, AiS allows the cell collective to adjust survival fate levels to react to environmental insults of different intensities. This mechanism is conserved across different epithelial cell lines, and might also contribute to mammary acinus morphogenesis.

### Mechanism of ERK/Akt wave propagation

We find that the ERK/Akt wave propagation depends on canonical EGFR signaling (Fig. 4J). This does not involve free diffusion of an EGFR ligand from apoptotic cells (Fig. 4C-E), but rather MMP-mediated cleavage of membrane-bound, precursor EGFR ligands such as amphiregulin (Dong et al., 1999). This might generate a trigger wave, in which each cell layer sequentially activates the next one through EGFR/MMP signaling. We show that ADAM17 might be required for shedding of the EGFR ligand from the apoptotic cell (Fig.4J). However, ADAM17 loss does not completely abrogate ERK/Akt waves, suggesting the involvement of additional players. Similar EGFR/MMP-dependent ERK waves control myosin contractility to organize collective epithelial cell migration in MDCK cells (Aoki et al., 2017; Hino et al., 2020). This suggests that EGFR/MMP-dependent ERK/Akt waves are an evolutionary-conserved pathway that can spatially regulate collective cell migration and AiS-dependent EH.

Another factor that shapes the signaling wave is the non-linearity of the MAPK cascade (Birtwistle and Kolch, 2011). The Raf/MEK/ERK cascade exhibits ultrasensitivity and adaptation (Ryu et al., 2015; Santos et al., 2007; Sparta et al., 2015), producing short ERK pulses of full amplitude even at low EGFR input. The finding that the number of ERK pulsing cells decreases in the successive layers of the wave (Fig.1I) suggests that the EGFR/MAPK cascade operates at the threshold input required for production of all-or-nothing ERK responses. This is consistent with the minute amounts of EGFR-ligand that might be released by MMPs. Thus, the observed 3 layers reach of the signaling wave might in part stem from the architecture of the MAPK network. A similar network logic might produce Akt pulses.

### Fate decision regulation by ERK/Akt pulse frequency

EGF-stimulated and apoptosis-triggered ERK/Akt pulses are virtually indistinguishable when compared outside of their spatial context (Fig.1 A and S1D). We show that low versus higher ERK pulse frequencies correlate with apoptosis and survival respectively (Fig. 5A-C). EGF stimulation can further increase pulse frequency, leading to cell cycle entry (Albeck et al., 2013). This suggests that low, medium and high ERK/Akt pulse frequencies respectively correlate with apoptosis, survival, and proliferation fates. In native tissues, combinatorial control of ERK/Akt pulses triggered by apoptosis or other sources might fine tune EH by contributing to both survival/proliferation fates.

How does a cell interpret the ERK/Akt signaling frequency into these specific fates? Several known ERK or Akt substrates have been implicated in regulating survival, such as the Akt substrate BAD (Datta et al., 1997) or the ERK substrate BIM (Harada et al., 2004). Another fate decision mechanism might involve ERK/Akt-dependent control of transcriptional programs (Avraham and Yarden, 2011). In the case of ERK, this involves immediate early genes (IEGs) that produce unstable transcripts with lifetimes of 30-40 min, that in turn regulate the transcription of delayed early genes (DEGs) with a lifetime of 1-3 h. Thus, one digital ERK/Akt pulse might switch on gene expression that fluctuates on the AiS timescale, allowing the cell collective to reset its survival fate and to remain responsive to future stimuli. High signaling frequencies might then allow to control the proliferation fate by regulation of IEGs such as Fra-1 that can linearly integrate ERK activity over time (Gillies et al., 2017). Another function of Akt pulses might be the single-cell regulation of metabolic activities required for enforcing specific fate decisions (Hung et al., 2017). Finally, we speculate that the ERK pulses might both regulate myosin contractility on time scales of minutes (Aoki et al., 2017; Hino et al., 2020) and fate decisions such as survival on time scales of hours. Thus, the MAPK pathway might function in an integrative fashion by regulating processes at different timescales to coordinate both cytoskeletal and survival/proliferation responses during morphogenetic processes.

### Population-level AiS responses

Our experiments and computational simulations show that upon the induction of apoptosis, e.g. through starvation or cytotoxic drugs, the feedback embedded in AiS allows the epithelium to dynamically adapt to different apoptotic rates. This might help to mitigate acute spikes of apoptotic rates that potentially compromise epithelial integrity, and to reach a new steady-state rate of apoptosis compatible with the regenerative capability of the epithelial tissue. Our mechanistic explanation of AiS is consistent with numerous reports in which targeting EGFR-ERK/Akt sensitizes cancer cells to chemotherapeutic agents (Barbuti et al., 2019; Holt et al., 2012; Smolensky et al., 2017). This provides a rationale to test further targeted/cytotoxic combination therapies to obtain better therapeutic responses in cancer.

### Physiological relevance of AiS signaling

Further studies are needed to understand the role of AiS *in vivo*. ERK activity waves have been observed *in vivo* in mouse epithelia (Hiratsuka et al., 2015). Here the ERK activity waves correlate with G2/M cell-cycle progression, but it is conceivable that additional inputs, not present in our starved monolayer, control this specific fate decision. Valon and colleagues (Valon et al., 2020) reported apoptosis-triggered, EGFR-dependent ERK pulses during development of the Drosophila pupal notum. This might provide robustness against external perturbations during development. These similarities suggest that AiS is a highly conserved mechanism regulating epithelial integrity throughout the animal kingdom.

## Material and methods

### Cell lines

Wild-type human mammary epithelial cells MCF10A cells were a gift of J.S. Brugge, Harvard Medical School, Boston, MA. ADAM17 KO MCF10A cells expressing H2B-iRFP (no selection), ERK1-mRuby2 (Blast), ERK-KTR-mCerulean3 (Hygro) and their WT counterpart were a gift of T. Aikin and S. Regot, Johns Hopkins School of Medicine, Baltimore, MD (Aikin et al., 2020). MCF10A cells were cultured in growth medium composed by DMEM:F12 supplemented with 5% horse serum, 20 ng/ml recombinant human EGF (Peprotech), 10 μg/ml insulin (Sigma-Aldrich/Merck), 0.5 mg/ml hydrocortisone (Sigma-Aldrich/Merck), 200 U/ml penicillin and 200 μg/ml streptomycin. All the experiments were carried out in starvation medium composed by DMEM:F12 supplemented with 0.3% BSA (Sigma-Aldrich/Merck), 0.5 mg/ml hydrocortisone (Sigma-Aldrich/Merck), 200 U/ml penicillin and 200 μg/ml streptomycin. Growth factor and serum starvation experiments were executed by removing the growth medium, 2 washes in PBS and replacement with starvation medium. MDCK and NRK-52E expressing EKAREV-NLS were a gift from K. Aoki (Aoki et al., 2013, 2017). MDCK and NRK-52E cells were cultured in DMEM supplemented with 10% FBS, 200 U/ml penicillin and 200 μg/ml streptomycin.

### Compounds and proteins

AZD5363 (Selleck Chemicals), Batimastat (MedChem Express), BMS-754807 (MedChem Express), Doramapimod (MedChem Express), Doxorubicin (MedChem Express), Etoposide (MedChem Express), Gefitinib (Sigma-Aldrich/Merck), SCH772984 (MedChem Express), SP600125 (MedChem Express), Trametinib (Selleck Chemicals) and zVAD-FMK (UBPBio) were dissolved in DMSO and preserved at −20°C in small aliquots to avoid thaw-freeze cycles. Cetuximab was purchased from MedChem Express. EGF (Peprotech) and IGF-I (Peprotech) were dissolved in aqueous solution and preserved at −20°C.

### Plasmids

The stable nuclear marker H2B-miRFP703 was generated by fusing the coding DNA sequence (CDS) of Homo sapiens H2B clustered histone 11 (H2BC11) with the monomeric near-infrared fluorescent protein miRFP703 CDS (Shcherbakova et al., 2016). ERK-KTR-mTurquoiose2 and ERK-KTR-mRuby2 were generated by fusing the ERK Kinase Translocation Reporter (ERK-KTR) CDS (Regot et al., 2014) with mTurquoiose2 (Goedhart et al., 2012) and mRuby2 (Lam et al., 2012) CDSs, respectively. FoxO3a-mNeonGreen was generated by fusing the 1-1188 portion of the homo sapiens forkhead box O3 a (FoxO3a) CDS with mNeonGreen CDS, a green fluorescent protein derived by Branchiostoma lanceolatum (Shaner et al., 2013). H2B-miRFP703, ERK-KTR-mTurquoiose2, ERK-KTR-mRuby2, FoxO3a-mNeongreen and LifeAct-mCherry were cloned in the PiggyBac plasmids pMP-PB, pSB-HPB (gift of David Hacker, Lausanne, (Balasubramanian et al., 2016)), or pPB3.0.Blast, an improved PiggyBac plasmid generated in our lab. For stable DNA integration we transfected the PiggyBac plasmids together with the helper plasmid expressing the transposase (Yusa et al., 2011). Transfection of MCF10A cells was carried out with FuGene (Promega) according to the manufacturer protocol. Stable clones with the different biosensors/optogenetic tools were generated by antibiotic selection and image-based screening.

The MOMP optogenetic actuator Cry2(1–531).L348F.mCh.BAX.S184E and Tom20.CIB.GFP, originally designated as the OptoBAX 2.0 system, and here termed as OptoBAX, was generated by Robert M Hughes (Godwin et al., 2019). Lyn-cytoFGFR1-PHR-mCit, expressing myristoylated FGFR1 cytoplasmic region fused with PHR domain of cryptochrome2 and mCitrine, here defined as OptoFGFR, was a gift from Won Do Heo (Addgene plasmid # 59776) (Kim et al., 2014). Lyn-cytoFGFR1-PHR-mCit was subcloned in a lentiviral backbone for stable cell line generation. pPB3.0-PuroCRY2-cRAF-mCitrine-P2A-CIBN-KrasCT, here defined as OptoRAF, was generated as an improvement of the previously reported light-induced ERK activation system, gift from Kazuhiro Aoki (Aoki et al., 2017). Cloning was done in two steps. First, the CRY2-cRaf sequence was cut from the pCX4puro-CRY2-cRAF plasmid using EcoRI and NotI. mCitrine was PCR amplified from the OptoFGFR plasmid adding NotI and XhoI sites and digested. Both sequences were ligated in the pPB3.0-Puro opened with EcoRI and XhoI. The GSGP2A-CIBN-KRasCT sequence (synthesized by GENWIZ) was digested with BsrGI and AflII and ligated in the opened pPB3.0-Puro-CRY2-cRAF-mCitrine.

### Live imaging

MCF10A cells were seeded on 5 μg/ml Fibronectin (PanReac AppliChem) on 24 well 1.5 glass bottom plates (Cellvis) at 1×10^5^ cells/well density two days before the experiment. Imaging experiments were performed on an epifluorescence Eclipse Ti inverted fluorescence microscope (Nikon) controlled by NIS-Elements (Nikon) with a Plan Apo air 20× (NA 0.8) or a Plan Apo air 40× (NA 0.9) objectives. Laser-based autofocus was used throughout the experiments. Image acquisition was performed with an Andor Zyla 4.2 plus camera at a 16-bit depth. The following excitation and emission filters (Chroma) were used: far red: 640nm, ET705/72m; red: 555nm, ET652/60m; NeonGreen: 508nm, ET605/52; GFP: 470nm, ET525/36m; mTurquoise2: 440nm, HQ480/40.

### MCF10A acini

3D live imaging experiments with MCF10A acini and the accompanying image analysis were performed as described in (Ender et al., 2020). Single MCF10A cells were embedded in growth factor-reduced Matrigel (Corning) and kept in DMEM/F12 supplemented with 2% horse serum, 20 ng/ml recombinant human EGF, 0.5 mg/ml hydrocortisone, 10 μg/ml insulin, 200 U/ml penicillin and 200 μg/ml streptomycin, which induced the development of acini. After 3 days in culture, EGF, insulin and horse serum were removed from the medium. 25 mM Hepes was to the medium added before imaging. Images were acquired on an epifluorescence Eclipse Ti2 inverted fluorescence microscope (Nikon) equipped with a CSU-W1 spinning disk confocal system (Yokogawa) and a Plan Apo VC 60X water immersion objective (NA = 1.2). Image acquisition was done at 16 bit depth with a Prime 95B camera (Teledyne Photometrics). A temperature control system and gas mixer (both Life Imaging Services) were used throughout imaging, as well as laser-based autofocus.

3D segmentation of nuclei and extraction of cytosolic/nuclear ERK KTR fluorescence intensity levels were performed using an updated version of the open source LEVER software (Winter et al., 2016). The apoptotic event was manually identified and annotated.

### Optogenetic experiments

Cells expressing OptoBAX, OptoFGFR or OptoRAF were kept in the dark for at least 24h before the experiments and all preparatory microscope setup was carried out with red or green light (wavelength > 550 nm). In the case of OptoBAX, cells expressing H2B-miRFP703 and ERK-KTR-mTurquoise2 were transfected with OptoBAX plasmids the day before the experiment to obtain sparse transfected cells (< 1% of the total cell population) in the MCF10A epithelium. Cells expressing the OptoBAX system were identified through the expression of Cry2(1–531).L348F.mCh.BAX.S184E fusion protein that contains mCherry. Fields of view containing confluent monolayers and at max 1-2 OptoBAX cells were selected for the experiment. Cells were illuminated with whole field blue light at 440nm every two minutes for the entire duration of the experiment to both induce translocation of BAX to mitochondria and to image ERK-KTR-mTurquoise2. Additionally cells were illuminated at 508nm to visualize Tom20.CIB.GFP, 555nm for mCherry (BAX fusion protein) and 640nm H2B-miRFP703. In the cases of OptoFGFR or OptoRAF, cells expressing H2B-miRFP703 and ERK-KTR-mRuby2 were infected with the lentiviral vector expressing Lyn-cytoFGFR1-PHR-mCit or transfected with pPB3.0.PURO.CRY2.cRAF.mCitrine.P2A.CIBN.KrasCT plasmid to generate stably expressing clones. Cells were imaged at 555nm for mRuby2 and 640nm H2B-miRFP703, and stimulated with 488 nm blue light for 100 ms at 3 W/cm^2, when required by the experiment protocol. Stimulation experiments were controlled by NIS jobs.

### Image analysis

For nuclear segmentation, we used a random forest classifier based on different pixel features available in Ilastik software (Berg et al., 2019). For the training phase, we manually annotated nuclear (in 20-50 cells) and background pixels in 16-bit time-lapse TIF images of the nuclear channel (H2B-miRFP703). The probability map of nuclei was then exported as 32-bit TIF images to perform threshold-based segmentation using CellProfiler 3.0 (McQuin et al., 2018). Expansion of nuclear objects by a predefined number of pixels was used to identify the area corresponding to the cytoplasm on a cell-by-cell basis, as described previously (Pargett et al., 2017). Cytosol/nuclear ratio was calculated by dividing the average cytosolic pixel intensity by the average nuclear pixel intensity. For visualization and quality control, we created images of segmented nuclei color coded according to ERK-KTR C/N. Single-cell tracking was performed on nuclear centroids in MATLAB using μ-track 2.2.1 (Jaqaman et al., 2008). For FRET experiments with MDCK and NRK-52E expressing EKAREV-NLS we used a partially modified pipeline. We used the donor channel for pixel classification of nuclei in Ilastik. We then used CellProfiler 3.0 to correct uneven illumination, to segment nuclei from the Ilastik nuclear prediction, and to calculate the FRET ratio, where the FRET image is divided pixel-by-pixel by the Donor image. Single-cell tracking was performed on nuclear centroids in MATLAB using μ-track 2.2.1.

Apoptotic events were annotated manually based on nuclear shrinkage to set time to 0. This approach showed higher accuracy compared to any available computational or marker-based methods.

Holes in the epithelial layer were identified using Ilastik’s machine learning algorithm starting from the manual annotation of holes in a combined ERK-KTR-mTurquoiose2 FoxO3a-mNeonGreen channel. After segmentation of the hole probability, their total area in each frame and field of view was calculated with Fiji.

### Extrusion assay

MCF10A cells were seeded in 6 well plates (TPP Techno Plastic Products AG) at 4×10^5^ cells and grown for 3-4 days till reaching the homeostatic confluent state. The evening before, the wells were washed twice with DMEM:F12, 0.3% BSA, 0.5 mg/ml hydrocortisone, 200 U/ml penicillin and 200 μg/ml streptomycin. After 16h, suspended material, including cell debris, was collected, concentrated and stained for 10 min with Hoechst and CellMask orange (Thermo Fisher). The stained debris were put in between two coverslips and acquired with a widefield fluorescence microscope with a 20x objective. Particles were segmented with CellProfiler 3.0 and data analyzed with R.

### Microfluidic device fabrication and preparation

Microfluidic devices were prepared as described previously (Ryu et al., 2015). Polydimethylsiloxane (PDMS) polymer (Dow Corning) was mixed with the catalyzer in 10:1 ratio in a plastic beaker. A first layer of 4–5 g was poured on the master and then degassed in a desiccator before solidifying at 80°C for 1 h. Eight‐well reservoir strips (Evergreen) were divided in 2 and then glued on the first layer using PDMS and solidifying at 80°C for 30 min. Finally, the second layer of 15–20 g of PDMS is used to finalize the device. The PDMS replica was cut and punched at the appropriate inlets and outlets. Plasma treatment was used to bond the PDMS replica to the 50 × 70 mm coverslip (Matsunami, Japan) to allow for proper sealing that resists the high‐pressure applied during the experiments. To enhance the bonding strength, the device was heated for 15 min in an 80°C dry oven. After bonding, the device was immediately filled by adding 200 μl of 10 μg/ml fibronectin solution in PBS to each outlet reservoir and put at 37°C. To increase coating efficiency in the device, 10 μl of the fibronectin solution was aspirated 3× from the cell seeding port after 1 h each before seeding cells.

MCF10A cell suspensions were prepared at a concentration of 1 × 10^7^ cells/ml. 30 μl of this cell suspension was added to the outlet reservoir and aspirated with a pipette from the cell reservoir inlet port. 250 μl complete medium was added to each outlet reservoir and 150 μl of starving medium to each inlet reservoir. Devices were kept overnight in the incubator. Before experiments, medium in the inlet and outlet reservoir was removed and 240 μl of appropriate medium was added to the inlet reservoir. Before imaging, devices were connected to an Elevsys OB-1 flow controller and the desired air pressure (0 or 500 mPsi was applied).

### Data analysis

To analyze and visualize single-cell ERK/Akt activity time series, we used Time Course Inspector (Dobrzyński et al., 2019) and we wrote a set of custom R/Python codes. All R notebooks with code required to reproduce the plots and the underlying data are available as supplementary information (link to online repository: http://dx.doi.org/10.17632/kd3n5k784m).

### Convolutional Neural Network (CNN)

The convolutional neural network (CNN) was trained as a binary classifier that discriminates between apoptotic and non-apoptotic cells based on their ERK activity time series. These come from four experiments where ERK activity in MCF10A cells was measured every minute. We manually annotated apoptotic cells by visual inspection of the movies and obtained 448 time series with a duration of 8h before the nuclear contraction. To create a balanced dataset and to avoid classification bias towards either of the classes, we selected 448 random time series of equal duration with cells that did not undergo apoptosis. We performed a data augmentation step by taking random 6h-long segments at every training epoch of the CNN analysis. We chose a plain CNN architecture as described previously (Zhou et al., 2016). It consists of several convolution layers followed by global average pooling (GAP) and a fully connected layer right before the output layer. The advantages of this architecture are a low number of parameters and a strong overfit counterweight with the GAP layer. The architecture comprises 6 successive 1D convolutional layers, all composed of 20 filters except for the last layer with 7 filters. In these layers, the convolution kernel sizes are (5, 5, 5, 5, 3, 3), respectively, with a stride of 1 and a padding that ensures a constant size of the input representation throughout the network. Each convolutional layer is followed by batch normalization and ReLU activation. The whole dataset (training and validation sets) was passed once through the network and the representation of time series after the GAP layer was projected in 2D with tSNE (Fig. 5A). Time series that maximized the model confidence for a given class were reported as prototypes. Time series for which entropy of the classification output was maximal (*i.e.* the model returns nearly 50% confidence for both classes) were reported as low-confidence prototypes.

### Peak detection

The number of ERK activity peaks was calculated with an algorithm that detects local maxima in time series. A local maxima is defined as a point that exceeds the value of its neighbors by a threshold that we manually set for each dataset. The thresholds were: 1.5e-1 for MCF10A cells; 5.0e-2 for the MCF10A/ADAM17KO co-culture experiments; 5.0e-3 for MDCK cells; 5.0e-3 for NRK cells.

### Identification of collective events

To estimate spatial correlations in ERK activity, for every field of view and a time-lapse frame we calculated the local indicators of spatial association (Anselin, 1995) using the `lisa` function from the `ncf` package available for R. The function assumes the value of ERK-KTR C/N fluorescence intensity and X/Y positions of the nucleus’ centroid. It then outputs the Moran’s I autocorrelation coefficient for cells within a radius, which we set to 100px, and the permutation two-sided p-value for each observation based on 1000 resamples. After thresholding cells for the correlation > 0.5 and the p-value < 0.05 we applied the dbscan algorithm from the R package `dbscan` to identify clusters of collective ERK activity. We assumed that a collective event consisted of a minimum 3 cells with the maximum distance between cells of 100px. This procedure yielded individual collective events in each frame of the time lapse movie. We chose only those collective events where ERK-KTR C/N was above a threshold. To estimate the threshold, we performed k-means clustering with 2 centers on individual ERK-KTR C/N values from all collective events. The threshold was the midpoint between the two cluster centers. Movies of nuclei color-coded according to ERK-KTR C/N with collective events indicated with convex polygons were prepared with `ffmpeg` software from individual png files produced by `ggplot2` in R scripts.

### Analysis of cells’ history

We matched the X/Y/T coordinates of manually annotated apoptotic events to single-cell time series using the `distance_join` function from the `fuzzyjoin` package for R. Using the maximum euclidean distance of 10px we automatically matched ~95% of apoptotic events. For further analysis, we chose only time series longer than 6h. To plot the ordered heatmaps in Fig. 4E, we overlaid the instances of collective events onto C/N ERK-KTR time series of single cells that underwent apoptosis. As a negative control, for each apoptotic event, we chose a random cell from the same time frame that did not undergo apoptosis during the entire experiment. We only compared sections of time series that matched the lifetime of apoptotic cells. To calculate the average percentage of non-apoptotic cells affected by collective events during a time window before apoptosis of the corresponding apoptotic cell (Fig.5F), we repeated the selection of random non-apoptotic cells 1000 times.

### Mathematical model

The set of ODE equations was solved numerically using the function ode from the `deSolve` package in R:

Where: *C(t)* - the total number of cells, *P_1-4_(t)* - protection steps, *k_apo_* - the rate of apoptosis, *k_a_* - the rate

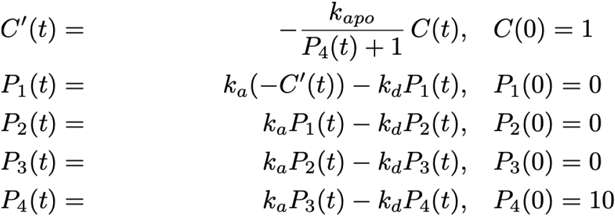

of protection activation, *k_d_* - the rate of protection deactivation. For all simulations *k_d_* was set to 1. Other parameters for scenarios in Fig. 4BC: (1) *k_apo_* = 0.01, *k_a_* = 5, (2) *k_apo_* = 0.05, *k_a_* = 5, (3) *k_apo_* = 0.05, *k_a_* = 2. The ratio *P_3_(t)/C(t)* was used as a measure of the average protection level per cell.

The R notebook and an interactive R/Shiny app are available as supplementary material (link to online repository: http://dx.doi.org/10.17632/kd3n5k784m). An interactive R/Shiny web-app to explore the parameters can be accessed online at https://macdobry.shinyapps.io/model-apo-prot/.

### Statistical analysis

All graphs were assembled and statistics were performed using R or Excel. The details of each statistical test performed are given in the legend accompanying each figure. Box plots depict the median and the 25th and 75th percentiles; whiskers correspond to maximum and minimum values in 1H, 1I, 1J, 2C, 2D, 7F, S2E, S2H, S3H, S4E, S5C, S5D or to 1.5 interquartile range drawn up to the largest observed point with the external observed points plotted as outliers in 4D, 4H, 4I, S1D, S4I, S4J, S4K, S5F, S6C. Histograms represent the mean ± standard deviation. XY line plots represent the mean and the shaded area represents the 95% confidence interval, except for 3A, 3C, S1E, S2I, S2J where it represents the standard deviation and S7A, S7I where it represents the standard error of the mean. Plots represent single-cell measurements or their distribution, except 6F, 6H, S6G-K that represent the distribution of biological replicates. The statistical tests used and the significance thresholds are given in each respective legend. N, when indicated, represents the N of apoptotic ERK/Akt waves analyzed.

## Acknowledgments

This work has been supported by grants from the Swiss National Science Foundation and the Swiss Cancer League to Olivier Pertz, from the Human Frontier Science Program to Olivier Pertz and Andrew Cohen, from ECU REDE (Research, Economic Development, and Engagement) and ECU THCAS (Thomas Harriot College of Arts and Sciences) to Robert M Hughes. We acknowledge support by the Microscopy Imaging Center at the University of Bern. We thank Sergi Regot and Tim Aikin for sharing ADAM17 KO cells, Kazuhiro Aoki for sharing the OptoRAF plasmid, MDCK and NRK-52E cells expressing EKAREV-NLS, and Won Do Heo for sharing the plasmids coding for OptoFGFR.

## Author contribution

PAG, MD and OP conceived the study. PAG and CD cloned and validated the biosensors. PAG performed experiments and image analysis. PE performed the 3D mammary acini experiment and ARC analyzed it. YB performed the microfluidic experiment. MD, M‐AJ and PAG analyzed the data. MD performed mathematical modeling. M‐AJ performed CNN analyses. CD built the optogenetic toolkit. RMH provided the OptoBAX tool and reviewed the article. OP supervised the work. PAG, MD, M‐ AJ, CD and OP wrote the paper.

