## Supplementary figures and legends for "Collective ERK/Akt activity waves orchestrate epithelial homeostasis by driving apoptosis-induced survival"

### Supplementary figure legends:

**Figure S1 (relative to main figure 1): Biosensors, image analysis pipeline and ERK/Akt wave propagation. (A)** Cartoon of the genetically-encoded ERK-KTR and FoxO3a biosensors. Both biosensors shuttle from nucleus to cytosol upon phosphorylation by ERK or Akt, respectively. A fused fluorescent protein (mTurquoise2 for ERK-KTR and mNeonGreen for FoxO3a) allows to visualize and quantify the shuttling. A genetically encoded stable nuclear marker H2B-miRFP703 is coexpressed with the two biosensors for nuclear segmentation and tracking. **(B)** Snapshots of MCF10A expressing the two biosensors and the nuclear marker, acquired in epifluorescence. **(C)** Schematic of the image analysis pipeline used in this study. A random forest algorithm implemented in Ilastik software (https://github.com/ilastik) is trained to produce nuclear-probability images from raw epifluorescence nuclear time-lapses. Nuclear probabilities are then segmented with CellProfiler (https://github.com/CellProfiler/CellProfiler) and tracked with u-track (https://github.com/DanuserLab/u-track). Nuclear segmented areas are expanded to generate a cytosolic ring. A single-cell ratio between the average cytosolic and the average nuclear pixel intensity is used as a readout of ERK or Akt activity. Final representations of ERK and Akt activities include: color-coded nuclear segmentations, single-cell time trajectories, and time heatmaps of C/N. Cells within heatmaps are clustered according to their relative time-averaged distance (*i.e.* adjacent trajectories in the heatmap correspond to neighboring cells). **(D)** Pulse amplitude of ERK and Akt activity in the presence or absence of EGF. Pulses were identified with the automatic peak detection described in the STAR methods section. Dots represent single pulses, box plots represent the quartile distribution. **(E)** Average (solid line) and standard deviation (shaded area) of ERK and Akt signaling trajectories of pulsing cells taken from different apoptotic events across neighborhood layers. The number of analyzed events: N=15 for the first layer of neighbors, N=16 for the second layer, and N=12 for the third layer. Black dashed line represents the time of nuclear shrinkage. Red, blue, and green dashed lines represent the peak time of ERK and Akt activity in the first, second, and third layers, respectively. **(F)** Distribution of the number of ERK-pulsing neighbors of 223 apoptotic events shown as the percentage of the total number of apoptotic events. Scale bars: (B, C) 50 μm.

**Figure S2 (relative to main figure 3): ERK/Akt activity wave is triggered by apoptotic events in a caspase-independent manner.** **(A)** Timelapse micrographs (left) and signaling trajectories (right) of apoptotic events induced by starvation, **(B)** 1 μM Doxorubicin or **(C)** 30 μM Etoposide in MCF10A cells expressing H2B-miRFP703, ERK-KTR-mTurquoise2 and FoxO3a-mNeonGreen. **(D)** Timelapse micrographs (left) and signaling trajectories (right) of an ERK/Akt activity wave induced by starvation in MCF10A cells treated with 100 μM zVAD-FMK to inhibit caspase activity. **(E)** Number of pulsing neighbors in ERK/Akt activity waves in presence or absence of zVAD-FMK. **(F)** MCF10A cells expressing H2B-miRFP703 and ERK-KTR-mRuby2 were transiently transfected with the OptoBAX optogenetic actuator. Snapshots of an apoptotic event induced by blue light illumination in the presence of zVAD-FMK. **(G)** Single-cell signaling trajectories from D. Highlighted trajectories correspond to cells inside the red contour in D. **(H)** The number of pulsing neighbors in ERK activity waves triggered by OptoBAX in the presence or absence of zVAD-FMK. **(I)** Nuclear area and **(J)** standard deviation of pixel intensity of the H2B-miRFP703 in the nuclear shrinkage and chromatin condensation phases in the presence or absence of zVAD-FMK, respectively. Solid line and shade represent the mean and standard deviation, respectively. **(K)** Fate of cells causing the ERK/Akt activity wave in the presence or absence of zVAD-FMK. Scale bars: (A, D) 50 µm.

**Figure S3 (relative to main figure 4): Apoptosis induced ERK/Akt wave depends on EGFR signaling. (A-F)** Timelapse micrographs (left) and signaling trajectories (right) of apoptotic events in cells expressing H2B-miRFP703, ERK-KTR-mTurquoise2 and FoxO3a-mNeonGreen and treated with (A) DMSO, (B) 1 µM Trametinib, (C) 5 µM AZD5363, (D) 1 µM Gefitinib, (E) 10 µg/ml Cetuximab and (F) 10 µM Batimastat. Snapshots show ERK and Akt activity in single-cells at different time points after nuclear shrinkage. Orange arrows indicate apoptotic cells. Highlighted trajectories correspond to cells showing ERK or Akt activity pulses after apoptosis. **(G)** Timelapse micrographs (left) and signaling trajectories (right) of MCF10A cells stably expressing H2B-miRFP703 and ERK-KTR-mTurquoise2 were transfected with two plasmids coding for two components of the OptoBAX optogenetic actuator. After illuminating cells with blue light, OptoBAX triggered apoptosis and a subsequent ERK activity wave. These examples show the effects of treatment with DMSO, 3 µM Gefitinib, 10 µg/ml Cetuximab or 10 µM Batimastat on ERK waves triggered by apoptosis. **(H)** Quantification of activated neighbors in different OptoBAX-induced apoptotic events in the presence or absence of 3 µM Gefitinib, 10 µg/ml Cetuximab or 10 µM Batimastat. Significance was obtained with a t-test with respect to DMSO treated cells. *, $P < 0.05$; ***, $P < 0.001$. **(I, J)** Timelapse micrographs (left) and signaling trajectories (right) of apoptotic events in cells expressing H2B-miRFP703, ERK-KTR-mTurquoise2 and FoxO3a-mNeonGreen and treated with (I) 10 µM Doramapimod or (J) 1 µM SP600125. Scale bar: 50 µm.

**Figure S4 (relative to main figure 4): Apoptosis induced ERK/Akt waves do not propagate through free diffusion and are partially dependent on ADAM17 protease. (A)** MCF10A cells expressing H2B-miRFP703, ERK-KTR-mTurquoise2 and FoxO3a-mNeonGreen were starved and treated with DMSO, 3 μM Gefitinib, 10 μM Batimastat or 10 μM BMS754807 and after 24 h stimulated with 10 ng/ml EGF or 10 ng/ml IGF-I. Heatmaps show 20 randomly chosen trajectories from two 20x FOVs per condition. **(B)** Theoretical representation of distance as a function of time due to flow or free diffusion. The red curve represents the constant flow at 25 μm/s. The blue curve corresponds to the free diffusion approximation with a diffusion constant of 1.4E-6 cm^2/s (estimation for amphiregulin based on similarly sized proteins). **(C)** Snapshots of MCF10A cells expressing H2B-miRFP703, ERK-KTR-mTurquoise2 and FoxO3a-mNeonGreen seeded in the microfluidic device. Dashed lines indicate the microfluidic channel. **(D)** images of fluorescent beads superimposed with tracking (red) in the two applied flow conditions. **(E)** Distribution of the speed of fluorescent beads. **(F, G)** Timelapse micrographs (left) and signaling trajectories (right) of apoptotic events in cells expressing H2B-miRFP703, ERK-KTR-mTurquoise2 and FoxO3a-mNeonGreen subject to no flow (F) or constant flow generated by 500 mPsi. **(H)** Timelapse micrographs of apoptotic events in wild-type and ADAM17 KO MCF10A cells expressing H2B-iRFP and ERK-KTR-mCerulean3. Arrowheads indicate apoptotic cells. White and red contours indicate the external border of the ERK activity wave. **(I)** Distribution of the time of the pulse maximum of the first neighbor that shows an ERK activity pulse in several apoptotic events in wild-type vs ADAM17 KO cells. **(J)** Distribution of the number of activated neighbors in wild-type vs ADAM17 KO cells across multiple apoptotic events. **(K)** Distribution of the number of activated neighbors in wild-type and ADAM17 KO co-culture conditions across multiple apoptotic events. **(L)** Micrographs of wild-type and ADAM17 KO cells in coculture experiments. **(M)**

Distributions of single-cell pulse frequency of wild-type and ADAM17 KO cells for a range of binned co-culture proportions. The proportion of 1 corresponds to a pure population of the respective cell line. Scale bars: (C, L) 200 µm; (F, G, H) 50 µm.

**Figure S5: ERK/Akt waves are not necessary for cell extrusion. (A)** Workflow of the extrusion assay: cells are washed to remove all previous debris, after 16 h the debris is collected, stained with Hoechst and CellMask orange and acquired with an epifluorescence microscope. **(B)** Epifluorescence images of debris for cells kept in growth or starvation medium and in the presence of the caspase inhibitor zVAD-FMK at different concentrations. **(C)** Quantification of the experiment in B with automatic counting of debris. **(D)** Quantification of the extruded material after 16 h of growth factor and serum starvation in the presence of Trametinib, AZD5363, Gefitinib and Batimastat. **(E)** MCF10A cells undergoing apoptotic epithelial extrusion in the presence of inhibitors interfering with ERK/Akt wave propagation. zVAD-FMK was used as negative control for its ability to hamper the extrusion process. **(F)** Distribution of extrusion time for several apoptotic events in MDCK monolayers treated with different inhibitors. The extrusion time was defined as the time between the initial nuclear shrinkage and the nuclear movement in the out of focus plane. **(G)** Timelapse micrographs of apoptotic events in MCF10A cells expressing H2B-miRFP703, ERK-KTR-mTurquoise2 and LifeAct-mCherry and treated with inhibitors, as indicated. Images were acquired with spinning-disk confocal microscope at two focal planes, one at the basal side to visualize the basal F-actin ring, and one medial to calculate the ERK-KTR C/N ratio. Arrows indicate the apoptotic cell. The red contours highlight the ERK activity wave triggered by apoptosis. Scale bar: 20 µm

**Figure S6 (relative to main figures 5 and 6): Supplementary data for apoptosis-induced and optogenetic-induced survival.** **(A)** Distribution of ERK peak frequency in apoptotic vs non-apoptotic MDCK and NRK-52E cells in the 6 h preceding apoptosis. The significance level calculated with the Wilcoxon test. **(B)** Workflow for the automatic quantification of collective events. Single-cell cytosol/nuclear ratio of the ERK-KTR channel is obtained as shown in S1C. In parallel, time of each apoptotic event is annotated on the base of nuclear shrinkage. Data of ERK dynamics were then used to automatically identify collective events and cross-referenced with apoptosis annotation. Scale bar: 100 μm. **(C)** The number of ERK pulses per hour in the period of 3 h preceding apoptosis compared to a period before that spans from the beginning of each track up to 3 h before apoptosis. **(D)** Percentage of secondary apoptotic events in the 2nd neighbors that received an ERK activity pulse during 4 h intervals after the primary apoptotic event. Error bars represent 95% CI. Dashed line and shaded grey area represent the percentage of pulsing cells in all 2nd neighbors and 95% CI. Significance level with respect to “>4h” calculated with Chi-square test and corrected with the Holm-Bonferroni method. **(E)** Cumulative distribution function of the probability of secondary apoptotic events in the 1st and 2nd layer neighbors of primary apoptotic events. The orange line corresponds to secondary apoptotic events that received an ERK activity pulse from the primary apoptotic event, and the blue line to events that did not receive a pulse. **(F)** Frequency of apoptotic events in MCF10A expressing H2B-miRFP703, ERK-KTR-mRuby2 and OptoFGFR under various light stimulation schemes of 100 ms blue light pulses, given at different periodicities. **(G)** Proportion of cells entering apoptosis upon treatment with 1 μM SCH772984, 1 μM Trametinib or 3 μM AZD5363 and simultaneous stimulation with OptoFGFR at 2 pulses per hour. Bars represent means and standard deviations from three independent experiments. Significance level was calculated by a t-test with respect to light-stimulated DMSO-treated condition. **(H-K)** Proportion of OptoRAF MCF10A cells undergoing apoptosis in the first 24 h after starvation in the absence/presence of OptoRAF light pulsing and with the following treatments: (H) 3 μM AZD5363, (I) 1 μM Trametinib, (J) 1 μM SCH772984 and (K) 10 μM Batimastat. Significance level was calculated with a t-test in 4 FOVs with respect to the DMSO unstimulated condition. *, P < 0.05; **, P < 0.01; ***, P < 0.01.

**Figure S7 (relative to main figure 7): AiS mediates steady-state apoptotic rates in doxorubicin treated MCF10A monolayers. (A)** Average number (normalized on 0 to 5 h) of living cells in four fields of view per condition in the same experiment. Shades represent SEM. **(B)** Apoptosis rate per hour, expressed as percentage of the initial population, of starved MCF10A monolayers treated with different concentrations of Doxorubicin. **(C)** Average ERK and Akt activity (solid lines) in the entire cell population of the experiment in A. Shades represent 95% CI. **(D)** Heatmaps of randomly-selected ERK and Akt activity trajectories in MCF10A monolayers treated with the different concentrations of Doxorubicin. **(E)** Apoptotic rate of MCF10A cells treated with 1.25 μM of doxorubicin compared to untreated cells under constant perfusion with the microfluidic device. **(F)** Average ERK and **(G)** Akt activity of cells treated with 1.25 μM of doxorubicin under constant perfusion. **(H)** Heatmaps of randomly selected single-cell trajectories from F and G. **(I)** Average number of cells per field of view of DMSO- or Doxorubicin-treated MCF10A monolayers in combination with inhibitors (10 μg/ml Cetuximab, 3 μM Gefitinib, 10 μM Batimastat, 1 μM Trametinib or 5 μM AZD5363). Shades represent SEM. **(J)** Apoptotic rate per hour in cells treated with Doxorubicin plus DMSO or other inhibitors as indicated. The dashed line represents a 3% apoptotic rate. This condition was identified as a steady-state apoptosis regime sustained by AiS.

### Video legends:

**Video S1 (Relative to figure 1D): collective ERK/Akt activity event triggered by apoptosis.** A collective ERK/Akt activity event shown in the H2B-miRFP703, ERK-KTR-mTurquoise2, FoxO3a-mNeonGreen, cytosol/nuclear segmentation, ERK and Akt activity channels. The cell highlighted with an arrowhead (cell no. 63 in the cytosol/nuclear segmentation channel) undergoes the apoptotic program, showing nuclear shrinkage at 0 min and nuclear fragmentation and blebbing about 1 h later. Immediately after the nuclear shrinkage, neighbors show a transient pulse of ERK/Akt activity that is then sequentially communicated to neighbors. The neighbors' topology relative to this event is shown in figure 1E. ERK and Akt signaling trajectories are shown in 1F.

**Video S2 (Relative to figure 2A-B): ERK activity waves triggered by apoptosis in MDCK and NRK-52E.** Examples of two apoptotic events in MDCK and NRK-52E expressing the ERK biosensor EKAREV-NLS (a gift from Kazuhiro Aoki). The two apoptotic cells are in the center of each window. Time 0 min corresponds to nuclear shrinkage. ERK and Akt signaling trajectories are shown in 2A.

**Video S3 (Relative to figure 4C): ERK/Akt activity waves occur under constant flow.** MCF10A cells expressing H2B-miRFP703, ERK-KTR-mTurquoise2 and FoxO3a-mNeonGreen were seeded in a microfluidic device to provide constant medium flow, generated by applying air pressure on a medium reservoir, or no pressure, for the negative control. Flow speed was calculated using fluorescent beads (top movie). Apoptotic cells are highlighted with white arrowheads. The movie speed for beads' movement and cells is different.

**Video S4 (Relative to figure 5G): fate of 1st layer neighbors of an apoptotic cell.** Fate and future ERK signaling of 1st neighbors of a primary apoptotic event (A) followed for > 1 day after the initial apoptosis. Three out of six first-layer neighbors (B, C and D) died in this time frame while the other three (E, F and G) survived. White circles highlight the apoptotic events, starting from nuclear contraction.

**Video S5 (Relative to figure 6B): ERK activity pulses induced by OptoFGFR and OptoRAF.** MCF10A cells expressing H2B-miRFP703, ERK-KTR-mRuby2 and the optogenetic actuators OptoFGFR or OptoRAF were stimulated with blue light (488 nm) for 100 ms at of 3 $W/cm^2$ intensity. The analysis of ERK dynamics shows that with both optogenetic actuators the ERK activation pulse is synchronous, transient, and homogenous.

**Video S6 (Relative to figure 6F): Frequency modulation of ERK pulses regulates the survival fate.** MCF10A cells expressing H2B-miRFP703, ERK-KTR-mRuby2 and the optogenetic actuators OptoFGFR were stimulated with pulsatile blue light (488 nm) for 100 ms with different interpulse periods: from 1 h to no pulsing. Images show nuclear segmentation color-coded according to ERK-KTR cytosol/nuclear ratio. A cyan blue bar above each image indicates the time of blue-light stimulation. Apoptotic events were annotated manually based on nuclear shrinkage and are represented in this video as white/red circles. This experiment shows that OptoFGFR pulsing every 1 to 3 h provides protection from apoptosis.

**Video S7 (Relative to figure 7G): Blocking AiS in the presence of acute apoptotic stress causes loss of epithelial integrity.** MCF10A expressing H2B-miRFP703, ERK-KTR-mTurquoise2 and FoxO3a-mNeonGreen were treated with 2.5 μM Doxorubicin together with DMSO, 10 μg/ml Cetuximab, 3 μM Gefitinib, 10 μM Batimastat, 1 μM Trametinib or 5 μM AZD5363. A machine learning approach was used to detect holes in the epithelial layer, here represented by black areas outlined in red. Inhibition of AiS determines a higher apoptotic rate that causes loss of epithelial integrity.

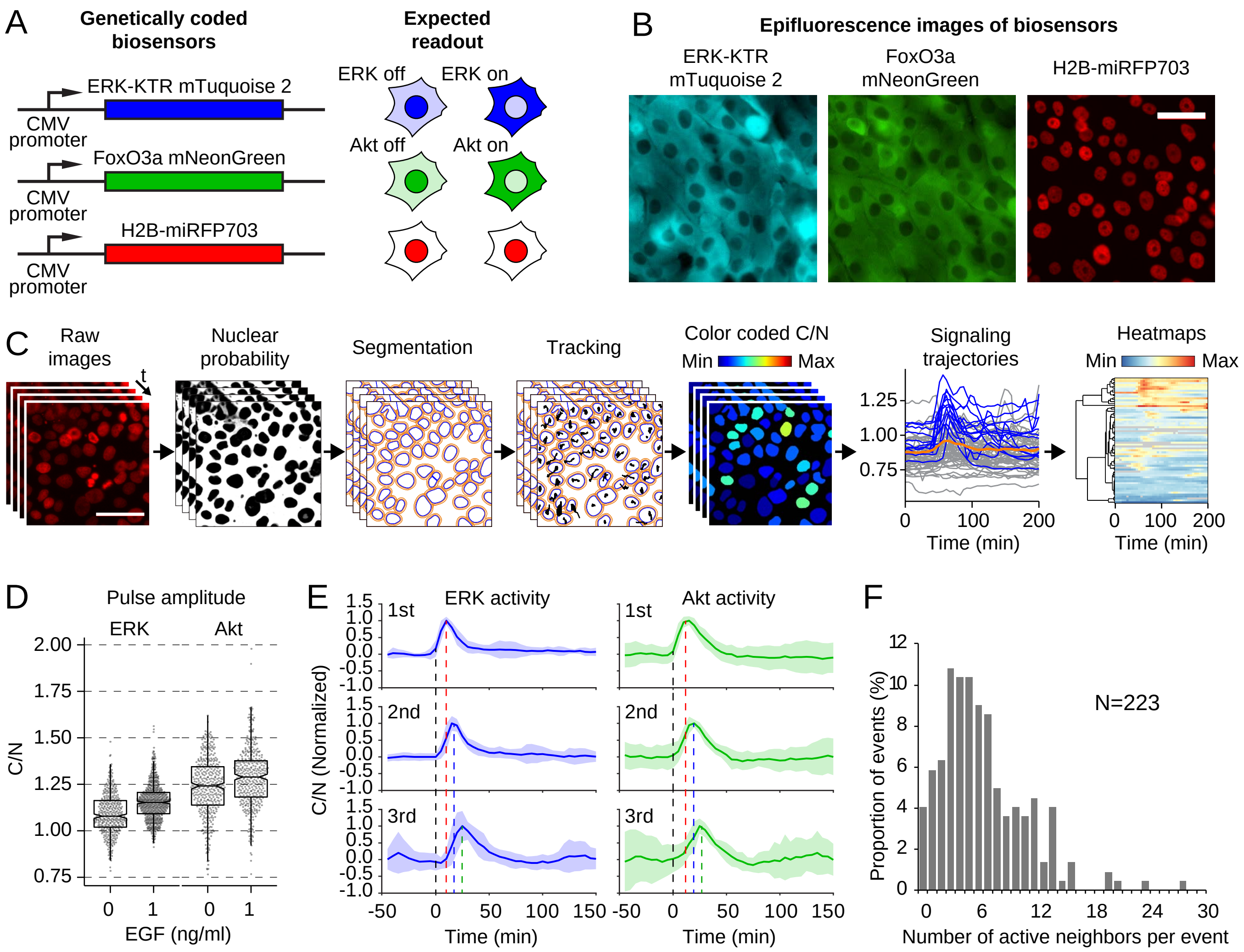


**Figure S1 (relative to main figure 1): Biosensors, image analysis pipeline and ERK/Akt wave propagation**

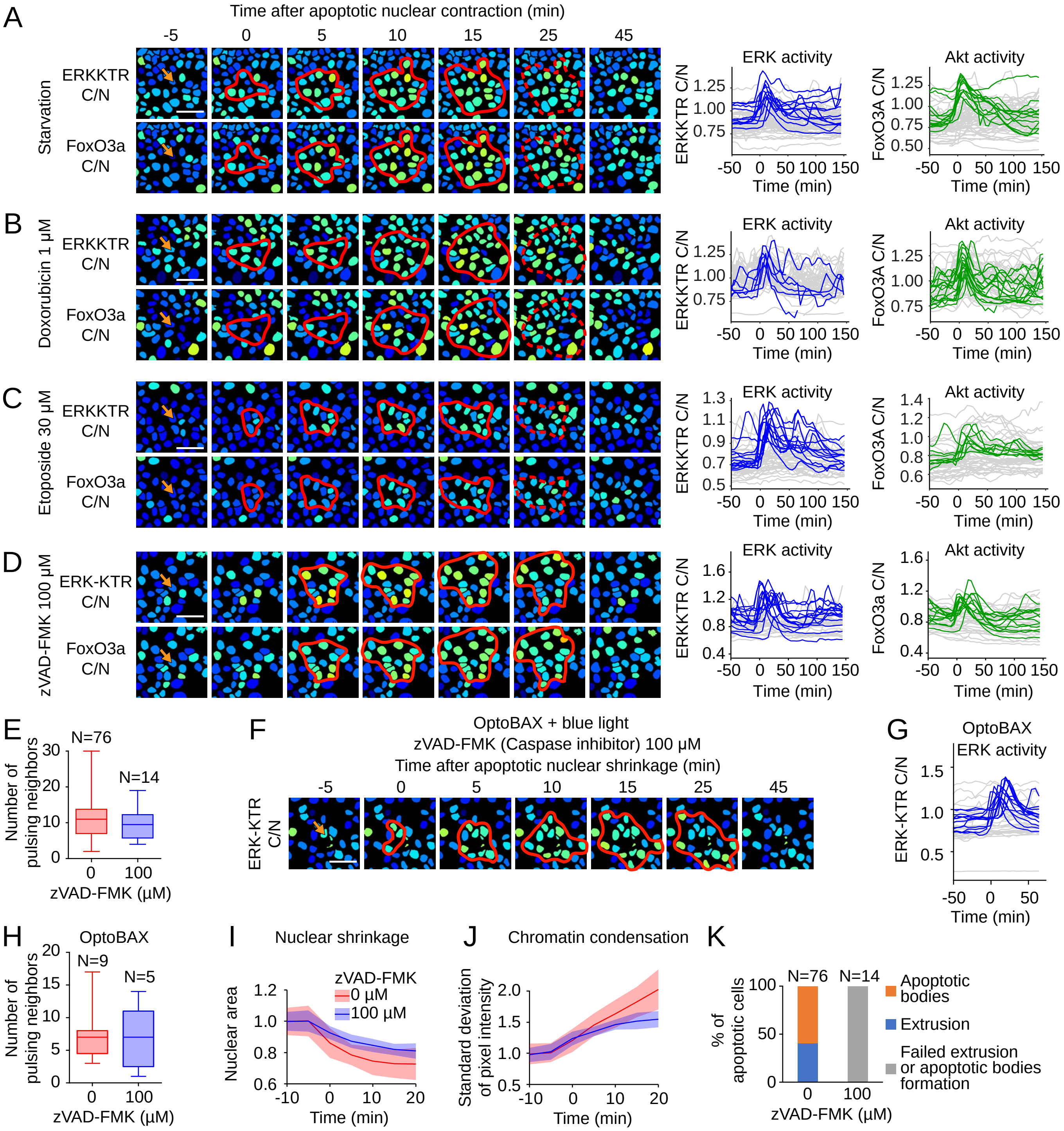


**Figure S2 (relative to main figure 3): ERK/Akt activity wave is triggered by apoptotic events in a caspase-independent manner.**

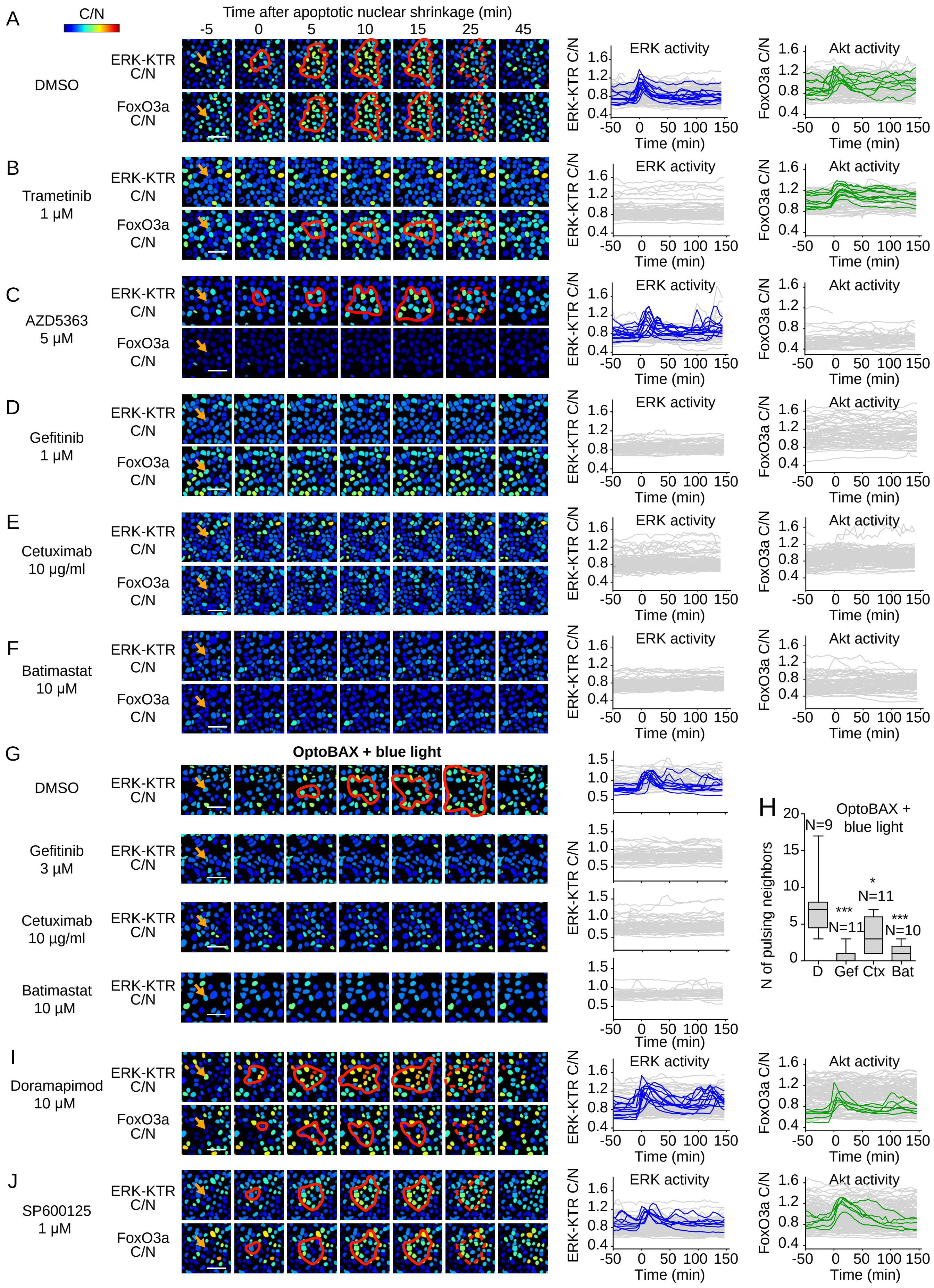


**Figure S3 (relative to main figure 4): Apoptosis induced ERK/Akt wave depends on EGFR signaling**

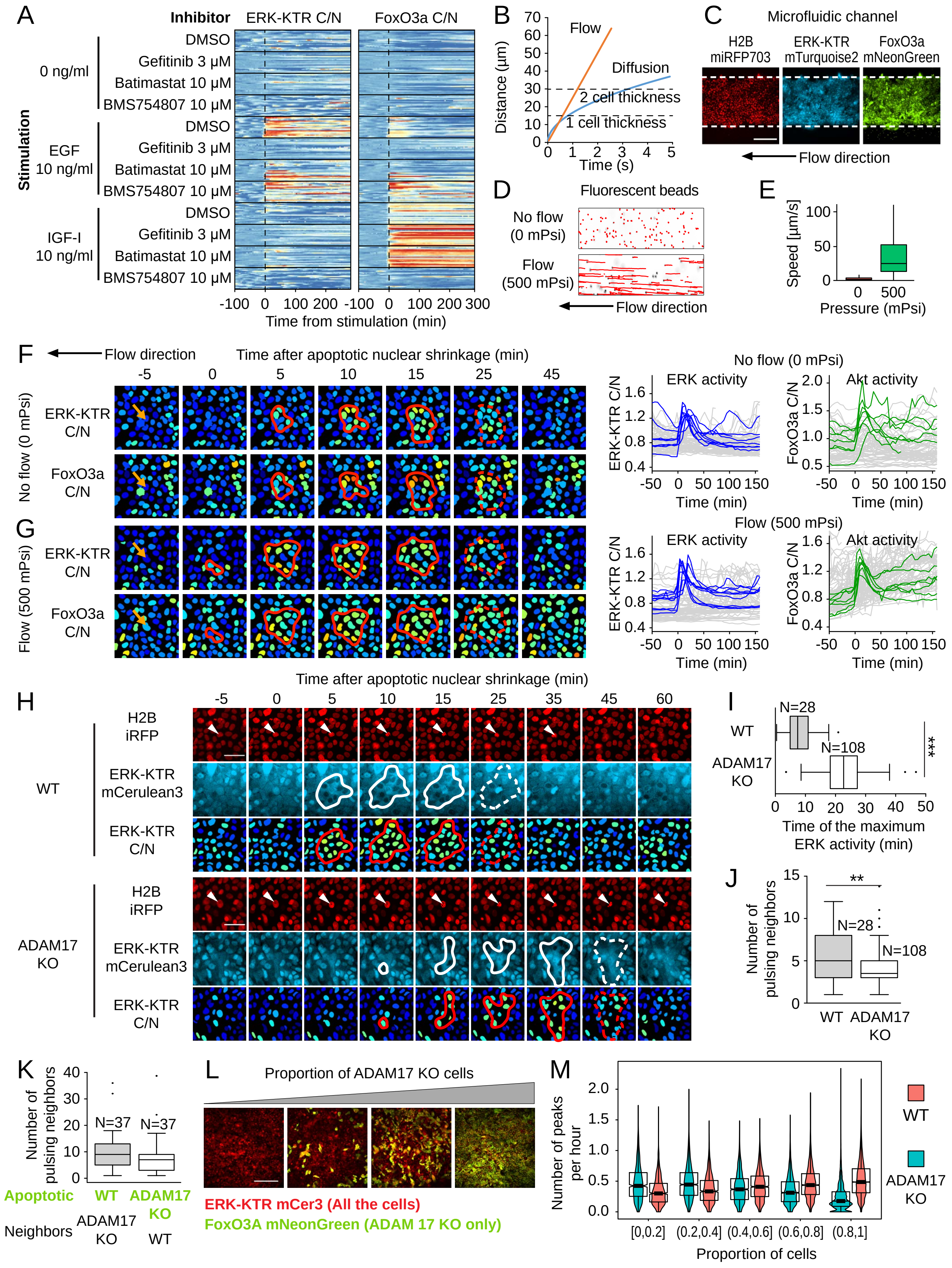


**Figure S4 (relative to main figure 4): Apoptosis induced ERK/Akt waves don't propagate through free diffusion and are partially dependent on ADAM17 protease**

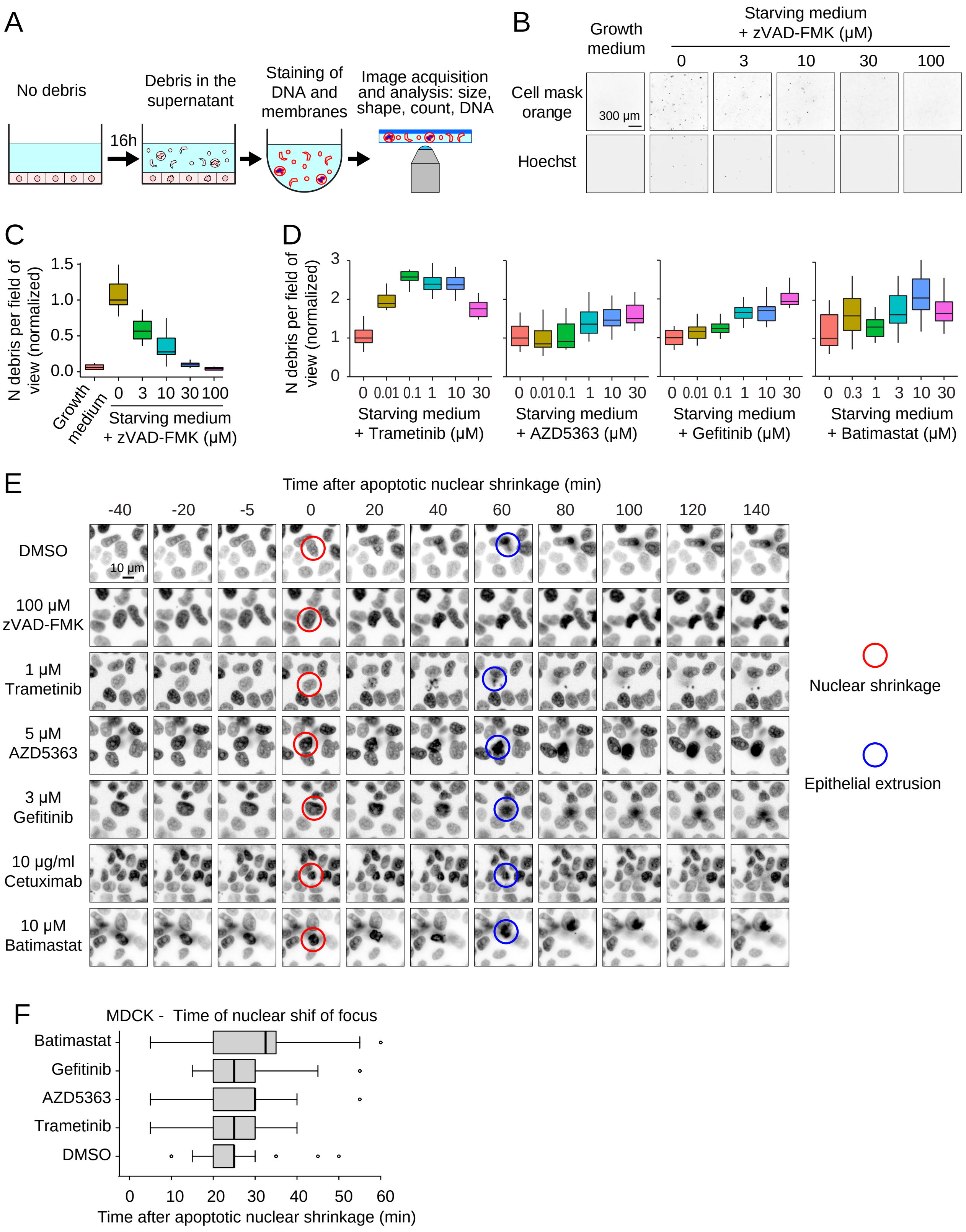


**Figure S5: ERK/Akt waves are not necessary for cell extrusion.**

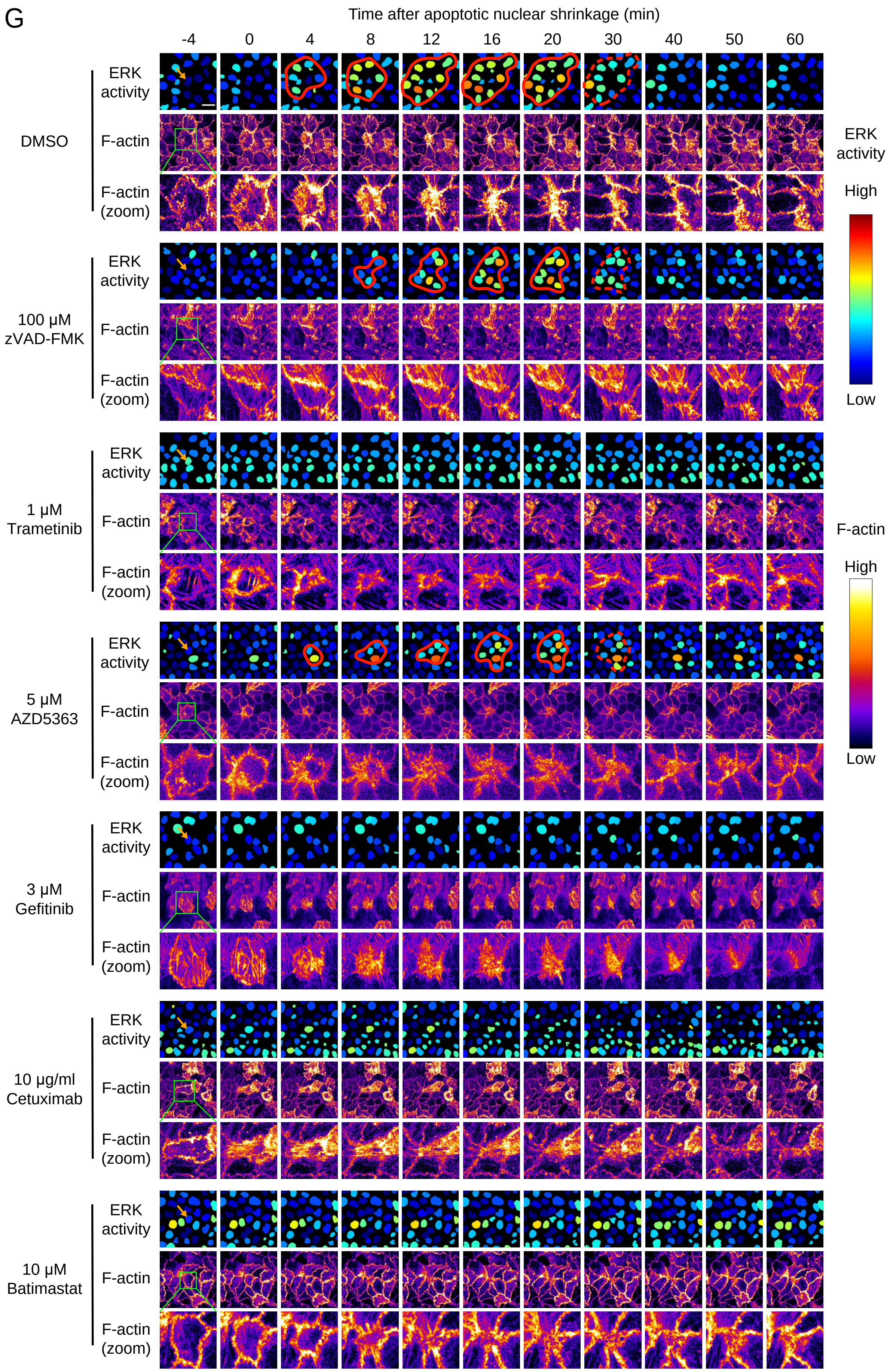


**Figure S5: ERK/Akt waves are not necessary for cell extrusion.**

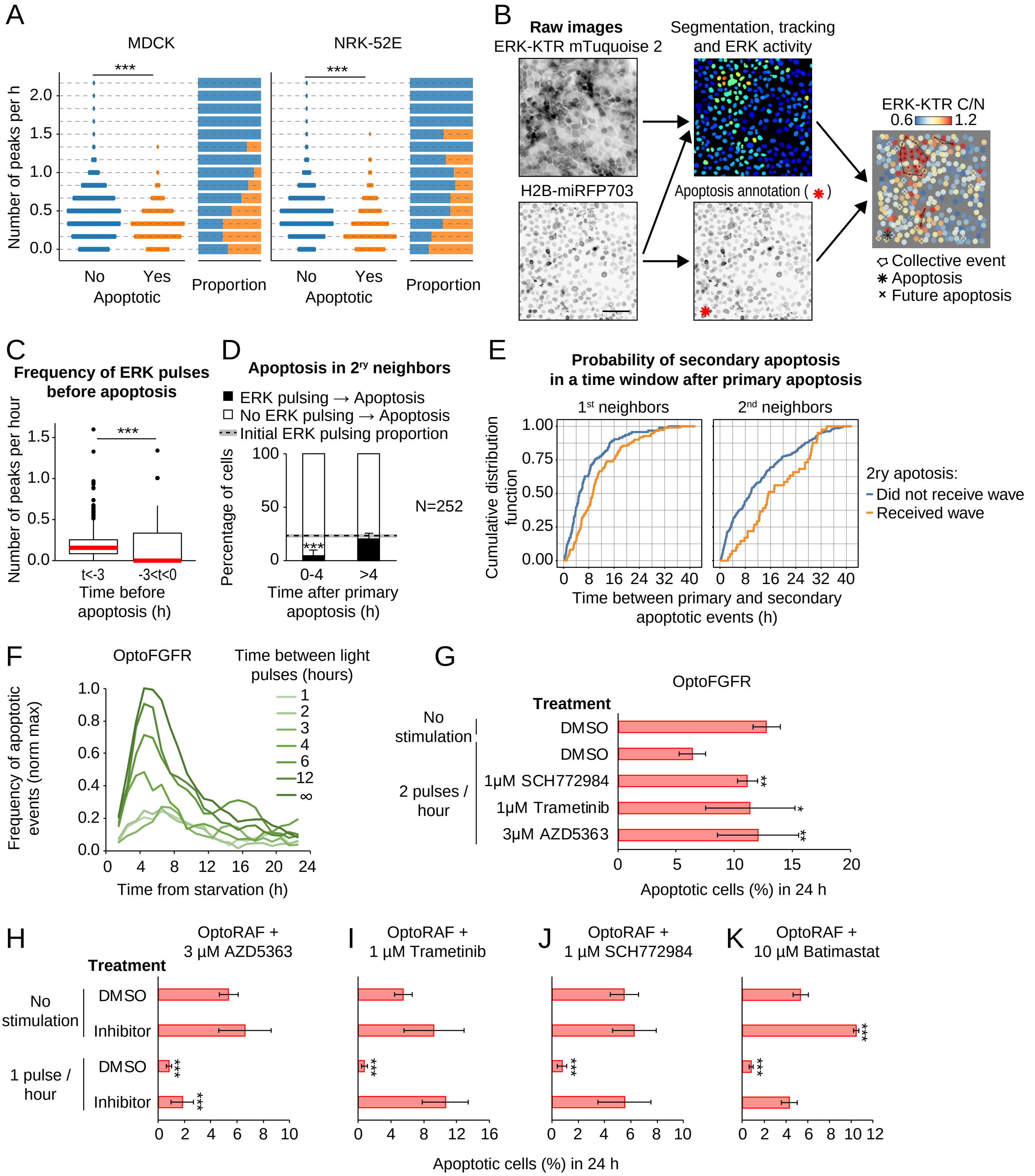


**Figure S6 (relative to main figures 5 and 6): Image and data analysis pipeline to identify collective events in signaling trajectories and optogenetically-induced survival.**

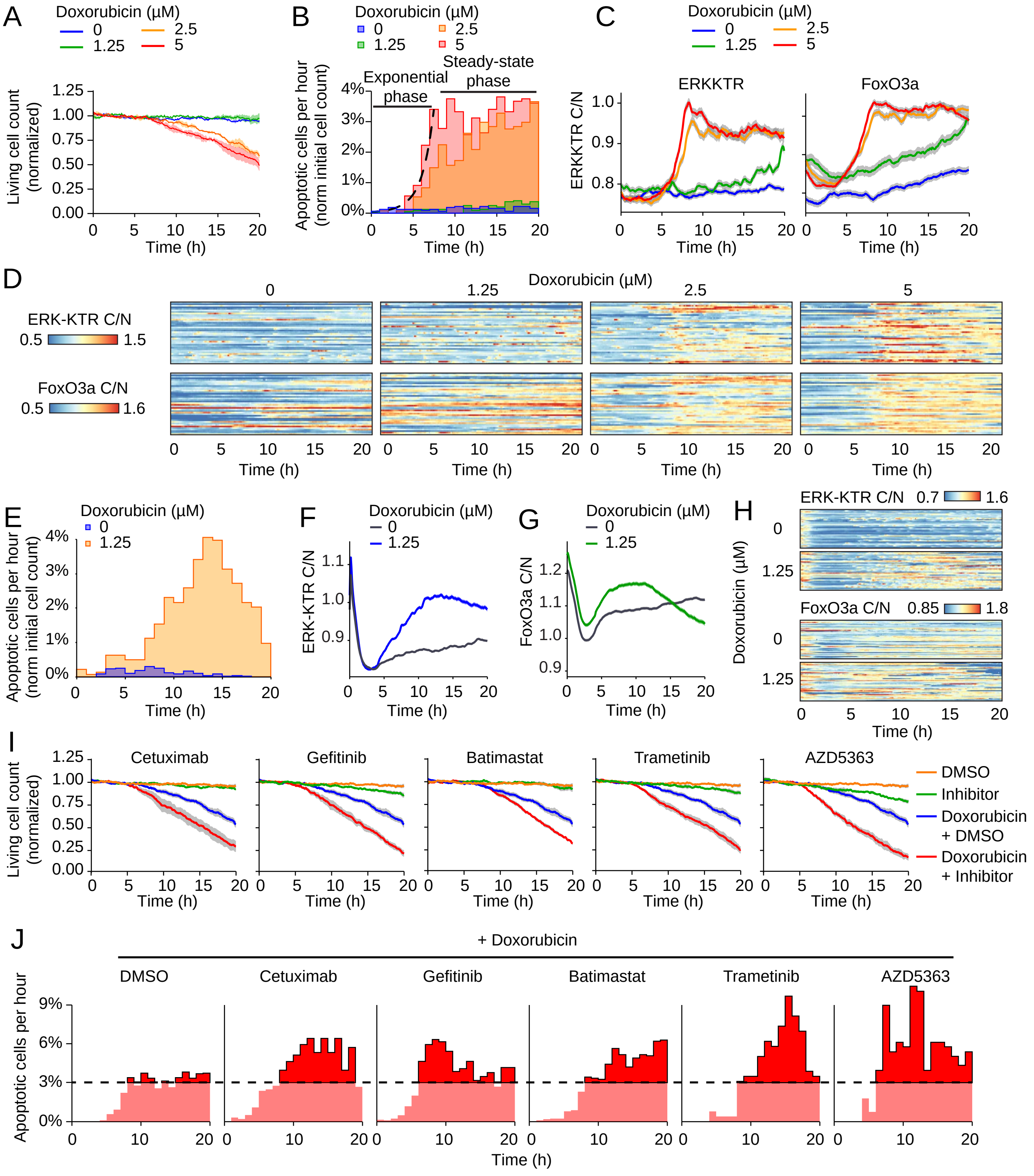


**Figure S7 (relative to main figure 7): AiS mediates steady-state apoptotic rates in doxorubicin treated MCF10A monolayers**
